# Flagellar beat state switching in microswimmers to select between positive and negative phototaxis

**DOI:** 10.1101/2023.12.20.572707

**Authors:** Alan C. H. Tsang, Ingmar H. Riedel-Kruse

**Affiliations:** Department of Mechanical Engineering, The University of Hong Kong, Hong Kong, China; Department of Molecular and Cellular Biology, and (by courtesy) Departments of Applied Mathematics, Physics, and Biomedical Engineering, University of Arizona, Tucson, USA

**Keywords:** phototaxis, adaptation, microswimmer, modeling, Euglena gracilis, two state switching, flagellar beating

## Abstract

Microorganisms have evolved various sensor-actuator circuits to respond to environmental stimuli. However, how a given circuit can select efficiently between positive vs. negative taxis under desired vs. undesired stimuli is poorly understood. Here, we investigate how the cellular mechanism by which the chiral microswimmer *Euglena gracilis* can select between positive vs. negative phototaxis under low vs. high light intensity conditions, respectively. We propose three general selection mechanisms for microswimmer phototaxis. A generic biophysical model demonstrates the effectiveness of all mechanisms, but which varies for each depending on specific conditions. Experiments reveal that only a ‘photoresponse inversion’ mechanism is compatible with *E. gracilis* phototaxis. Specifically, a light-intensity dependent transition on the sub-second time scale between two flagellar beat states responsible for forward swimming vs. sideway turning ultimately generates positive phototaxis at low light intensity via a run-and-tumble strategy and negative phototaxis at high light intensity via a helical klinotaxis strategy. More generally, a picture emerges where a variety of *E. gracilis* behaviors over a wide range of light intensities as reported in the literature can be explained by the coordinated switching between just these two flagellar beating states over time. These results provide design principles for simple two-state switching mechanisms in natural and synthetic microswimmers to operate under both noisy and saturated stimulus conditions.

**LAY ABSTRACT:** Our experimental and theoretical results explain how the single cell *Euglena gracilis* achieves both positive and negative phototaxis. Our insights are then able to synthesise a larger number of previously described observations on *E. gracilis* photoresponses and photobehaviors due to a concise two-state model of flagellar beating. These insight will likely inform the behaviors of other natural microswimmers as well as the design of synthetic ones.

## INTRODUCTION

Natural microswimmers perform versatile feedback control strategies by selecting cell motility behaviors in response to environmental stimuli [1–11]: Bacteria alternate between running and tumbling states to navigate chemical gradients [3–6]. Chlamydomonas reinhardtii coordinate their cis and trans flagellum to transition between positive and negative phototaxis [12–16]. Volvox carteri control their positive and negative phototaxis by accelerating or ceasing their anterior flagella [17, 18]. Actually, many organisms of different body size and flagella numbers are currently studied, promising the understanding of generalizable biophysical laws as well as a versatile solution spectrum to ‘navigation problems’ [19, 20]. How many of these microswimmers develop feedback response mechanisms to select between different motility behaviors, e.g., changing from positive to negative taxis, is not well understood.

*Euglena gracilis* is an ideal model to study these questions [9, 11, 21, 22]. These single celled organisms can perform positive and negative phototaxis at very low and high light intensities, respectively (i.e., from *<* 50 lux to *>* 10, 000 lux) (Fig. 1a,b) [9–11, 23]. *E. gracilis* rely on this transition to optimize photosynthesis, avoid photodamage by strong light, and regulate signal transductions that stimulate biological processes [9, 11, 24–27]. *E. gracilis* cells have an ellipsoidal shape with length and diameter of *∼* 50 *µ*m and *∼* 10 *µ*m, respectively, have one (or multiple) photosensors situated at or close to the ‘eyespot’ and have one main flagellum for propulsian and steering that beats with 20-40 Hz [11, 28, 29]. They swim in direction of their long-axis at *∼* 50 *µ*m/s while rolling around this long axis at a frequency of *ω≈* 1 Hz [21, 30, 31] (Fig. 1c). *E. gracilis* are ‘pullers’ [2], i.e., the flagellum is situated at the front of the cell from the perspective of the swimming direction (Fig. 1c). Multiple photoreceptors (e.g., flavins, PAC (Photoactivated adenylyl cyclase)) and various signaling pathway components between receptor and flagellar actuation have been identified, yet many open questions regarding these sensors remain as well [11, 32, 33].

**Figure 1.**
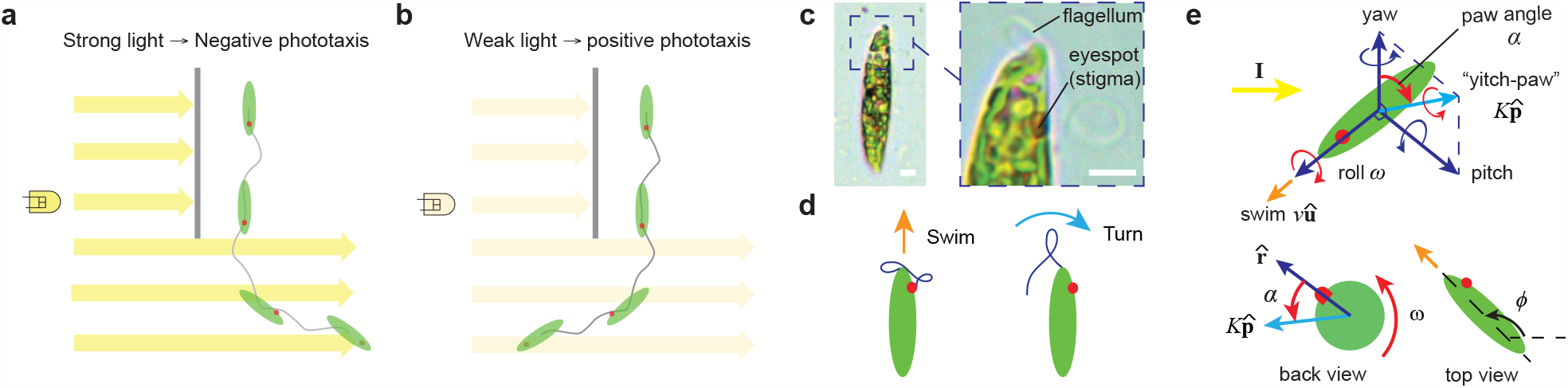
How microswimmers like *Euglena gracilis* achieve the selection between negative and positive phototaxis for different light intensities is unclear. **(a)** Schematic of *E. gracilis* performing negative phototaxis to swim away from strong light intensity. **(b)** Schematic of *E. gracilis* performing positive phototaxis to swim towards weak light intensity. **(c)** *E. gracilis* has a red eyespot (stigma) that shades its photoreceptor and a flagellum responsible for motility; scale bars: 5 *µ*m. **(d)** *E. gracilis* has two main beat patterns that primarily achieve forward swimming and sideward turning behavior [21]. **(e)** Schematic of a biophysical feedback model between *E. gracilis’s* photo sensor and flagellum established previously that allows for a rich set of photore-sponsive behaviors [21] (see text and Supplementary Material for model parameters and additional details).

*E. gracilis* has two distinct flagellar beating states that are responsible for forward swimming and sideways turning, respectively (Fig. 1d) [21, 22, 34, 35]. The photoresponsive actuation of both states is controlled by a feedback between the photosensor and the flagellum (Fig. 1c-e), which can lead to a significant number of different photo-responsive behaviors *E. gracilis* [9, 11, 21, 21, 22, 24, 25, 30, 31, 36, 37], and among those are positive and negative phototaxis (Fig. 1a,b). Furthermore, *E. gracilis* shows ‘step-down response’ vs. ‘step-up response’ to a decrease vs. an increase in light intensity at a low vs. high background light intensity, respectively [11, 22, 37], and both responses correspond to transitions from the flagellar beating state for swimming to the state for turning (Fig. 1d), and which will typically relax back to swimming after stimulus reversal or a longer adaptaion time [21, 22]. The phototaxis response of chiral microswimmers can then be modeled with respect of the three vectors 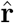, û, and 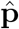 (Fig. 1e) [21].

In this paper, we now focus on the long standing questions on how *E. gracilis* achieves positive and negative phototaxis at low and high light intensities (Fig. 1a,b), respectively, and how a cell can select between both behaviors depending on overall light conditions [9, 11]. We take a fresh approach by using high-speed imaging and modeling. We propose three general mechanisms capturing phototaxis transition in microswimmers and verify the relevant mechanism(s) for *E. gracilis* by comparing our theoretical results with our experiments. We investigate flagellar beat responses upon step-ups or step-downs in light intensity at different overall light intensity levels, and how the cell then can achieve a set of different phototaxis strategies. We ultimately identify a single beat coordination mechanism for *E. gracilis* that explains a significant variety of short and long term photobehaviors and search strategies over a wide range of light intensities, and which in particular suggest that positive vs. negative phototaxis are achieved via a biased random walk vs helical klinotaxis, respectively [38, 39].

### Background

Using high-speed microscopy, we recently described in fine detail two distinct flagellar beating states that are responsible for forward swimming and sideways turning, respectively (Fig. 1d) [21]. The existence of two beating states had actually been noticed in the literature for a long time [22, 34, 35] - although without measuring the flagellar waveform sequence in detail given the limited imaging technology available at the time - instead papers usually just focused on the resulting overall cell movement [22]. The one exception we found was Diehn 1975 [40], who reported the switching between both beating states for the case of negative photoxis using high speed microscopy (at 100 and 125 Hz); this paper also stated that positive phototaxis could not be studied equivalently as the required low light levels prevented high speed imaging. Furthermore, neither this nor any of the following papers traced out a shape sequence for either of these beating states. Only in 2017, Rossi et al. [41] reported on the forward swimming beat pattern waveform in the absence of changing light stimuli, followed then by out work reporting on both beat patterns in high detail under various light stimuli [21]. Hence, we now have the high speed imaging and analysis technology to revisit many questions on photobeahviors (and behaviors in responses to other stimuli) in *E. gracilis* [9, 11].

Previous work on *E. gracilis* phototaxis has been mostly focused on the statistics of the swimming directions under different light stimuli, little is known about the flagellar mechanics and beat patterns that lead to these various photobehaviors [10, 11, 25]. In particular, it is not well understood how *E. gracilis* achieves positive and negative phototaxis at low and high light intensities (Fig. 1a,b), respectively, and how a cell can select between both behaviors depending on overall light conditions [9, 11]. A significant number of different photo-responsive behaviors have been reported for *E. gracilis*, e.g., positive and negative phototaxis [9, 11, 24, 25], polygonal swimming motion [21], localized spinning [21], ordered motion in polarized light [36], avoidance turning when encountering a light barrier [21, 30, 31], and a ‘step-down response’ vs. ‘step-up response’ [11, 22, 37]. In particular, the latter two represent in both cases an abrupt transition from the cell swimming forward to the cell tumbling around its axis after the light intensity has changed, and where the ‘step-down’ vs. ‘step-up’ refer to a decrease vs. an increase in light intensity at a low vs. high background light intensity, respectively; the cell then typically returns back to the swimming state upon stimulus reversal or after a longer adaptation time [11, 37]. Both step responses correspond to a transition from the flagellar beating state for swimming to the state for turning (Fig. 1d) [21, 22].

It has been proposed that positive and negative phototaxis can be generated by the frequent actuation (and relaxation) of these step responses: as the light intensity detected by one or multiple photosensors varies over time, they cell switches back forth between turning and forward swimming while the cells reorients itself relative to the light source [9]. Here cellular structures like the stigma or the green organelles (Fig. 1c) are supposed to partially shade these photosensors depending the light direction relative to the cell body orientation (Fig. 1e) [9, 42]. In contrast, multiple authors claim to have found experimental evidence that rejects this mechanism [22, 43], for example, given how cells respond to polarized light or multiple simultaneous light sources [44–46], or since allegedly no directional positive phototaxis was ever observed (and which is considered to be distinct from photoaccumulation) [47]. To support or refute these and other potential mechanisms underlying *E. gracilis* photobeaviors, various mathematical models have been investigated e.g., [21, 42, 48]. Independent of mechanism, the phototaxis response of chiral microswimmers can be modeled with respect of the three vectors 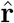, û, and 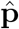(Fig. 1e) [21]: The direction and light intensity of a far away light source is denoted by **I**. A photoreceptor in the microswimmer senses this light signal **I**, and where 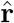 denotes the receptor’s direction of maximal sensitivity. This signal is converted into a specific flagellar beat pattern (Fig. 1d), which can affect the swimming speed *v* (‘surge’) along the long axis û, rolling frequency *ω* (‘roll’, positive for anticlockwise rotation) around û, and a more complex side-way turning (‘yitch-paw’) around 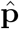 with strength *K* via modulation of a paw angle *α* in a light dependent manner. To compare with experimental results, we further denote the projection of the cell orientation in the 2D plane of the lab frame by the orientation angle *ϕ*; if not stated otherwise, the light vector **I** is parallel to this 2D plane. These motions then affect the orientation and position of the cell in 3D space, which affects the detected light, ultimately leading to a complex feedback (Fig. 1e) [21]. Overall, the understanding of a coherent mechanisms underlying this sensor-actuator feedback that generates all these various photobehaviors under various stimulus conditions is still lacking - both from a molecular signalling as well as a general information processing perspective.

## RESULTS

### Negative vs. positive phototaxis

First, we experimentally recorded *E. gracilis*’s swimming trajectories and population responses during positive and negative phototaxis (Fig. 2). White light stimuli were applied from the side of the observation chamber with background light from below (see supplements for full experimental methods). Negative phototaxis swimming trajectories were parallel to the light vector with slight oscillations due to cellular rotation (Fig. 2a,b; Supplementary Movie S1; background ~ 100 lux, stimulus *>* 2, 000 lux). Positive phototaxis trajectories appeared almost random with frequent turns of *∼* 5*−*10 seconds and a weak bias in orientation of *ϕ∼ π/*2 towards the light (Fig. 2c,d; Supplementary Movie S2; background *∼* 0 lux, stimulus *<* 50 lux; cells were imaged under red light which does not cause responses [23]). Over the time scales of 5 minutes the cells accumulated close to the light source (Fig. 2c, Supplementary Movie S2). The orientation distributions for negative phototaxis was much narrower than that for positive phototaxis, i.e., 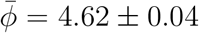 (mean *±* sem, throughout the paper if not stated otherwise) vs. 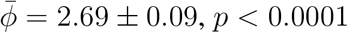 (Fig. 2b,d).

**Figure 2.**
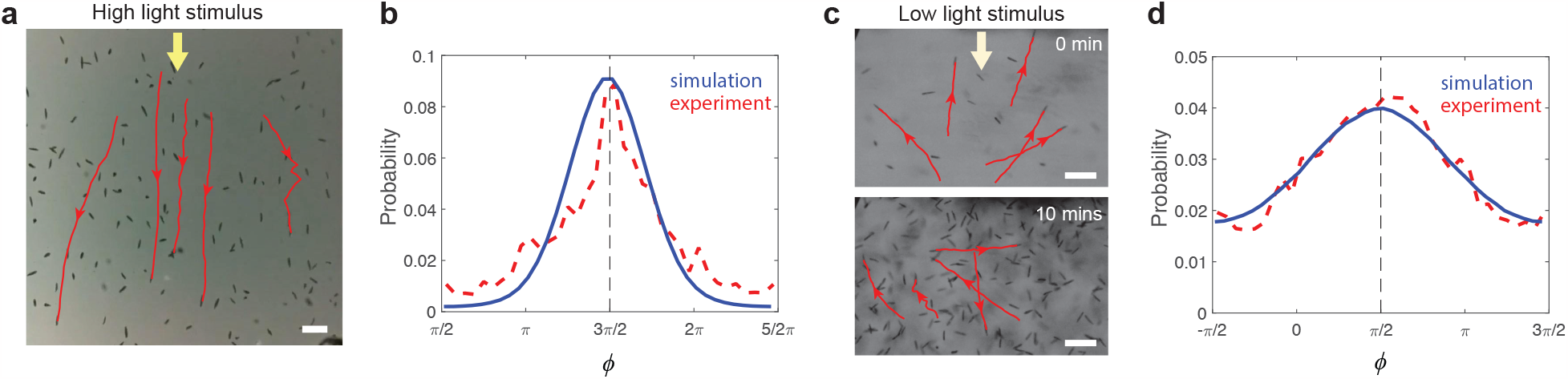
*Euglena gracilis* exhibits directional negative phototaxis and noisy positive photo-taxis. **(a)** During negative phototaxis, the cells show directed swimming away from a high light intensity srouce (Supplementary Movie S1). **(b)** Cell orientation *ϕ* during negative phototaxis shows directed swimming away from light, with a sharp peak in the light direction. **(c)** During positive phototaxis, the cells do not show directed swimming towards the light source, but slowly accumulate close to the light source over 10 minutes (Supplementary movie S2). **(d)** Cell orientations during positive phototaxis show a wide distribution, with a weak bias towards the light. The red lines in (b) and (d) depict the experimental results obtained from the tracking data in a 1 and 5 minute interval, respectively, with each cell tracked over 3 seconds. 406 and 1, 320 cells were tracked in experiments in (b) and (d), respectively. The blue lines in (b) and (d) depict the results obtained from Monte-Carlo type simulations of 1, 000 runs with duration of 600 time units. See Supplementary Section 3.9 and 4.2 for model parameters and additional details. Scale bars: (a) 100 *µ*m, (c) 100 *µ*m.

At medium light intensities (*∼* 100 lux), cells showed no phototaxis. Previously, we also reported on polygonal swimming (*∼* 500 *−* 1, 500 lux) and localized spinning (*>* 3, 000 lux) - both without observing any phototaxis [21]. The resulting statistical distributions are consistent with earlier reports (Fig. 2b,d) [10, 11, 25]. We conclude that negative vs. positive phototaxis appear to be achieved via directional steering (helical klinotaxis) vs. a biased random walk, respectively.

### Modeling possible phototaxis selection mechanisms

This data raises the question of what feature in the sensor-actuator circuit (Fig. 1e) changes when performing positive instead of negative phototaxis, i.e., operating in a very low instead of high light intensity environment, and we propose three possible ‘selection mechanisms’ (Fig. 3b-d): (1) Light-dependent angular turn: The paw angle *α*, which determines the orientation of the yitch-paw vector around which the cell turns upon detection of light (Fig. 3a), is changing, for example, it increases by *π* from *α* to *α*^*′*^ (Fig. 3b). (2) Photoresponse delay: The cell’s turning response upon detecting light is delayed, e.g., by half a roll cycle (Fig. 3c). (3) Photoresponse inversion: The effect of light stimuli onto the flagellar beating states for swimming and turning (Fig. 1d) are swapped, i.e., the cell frequently turns in the dark (step-down response) but swims straight when it detects light (step-up response) (Fig. 3d). Hence for all three mechanisms, the cell should be able to turn towards the light source instead of away from it, therefor leading to positive phototaxis at low light intensities.

**Figure 3.**
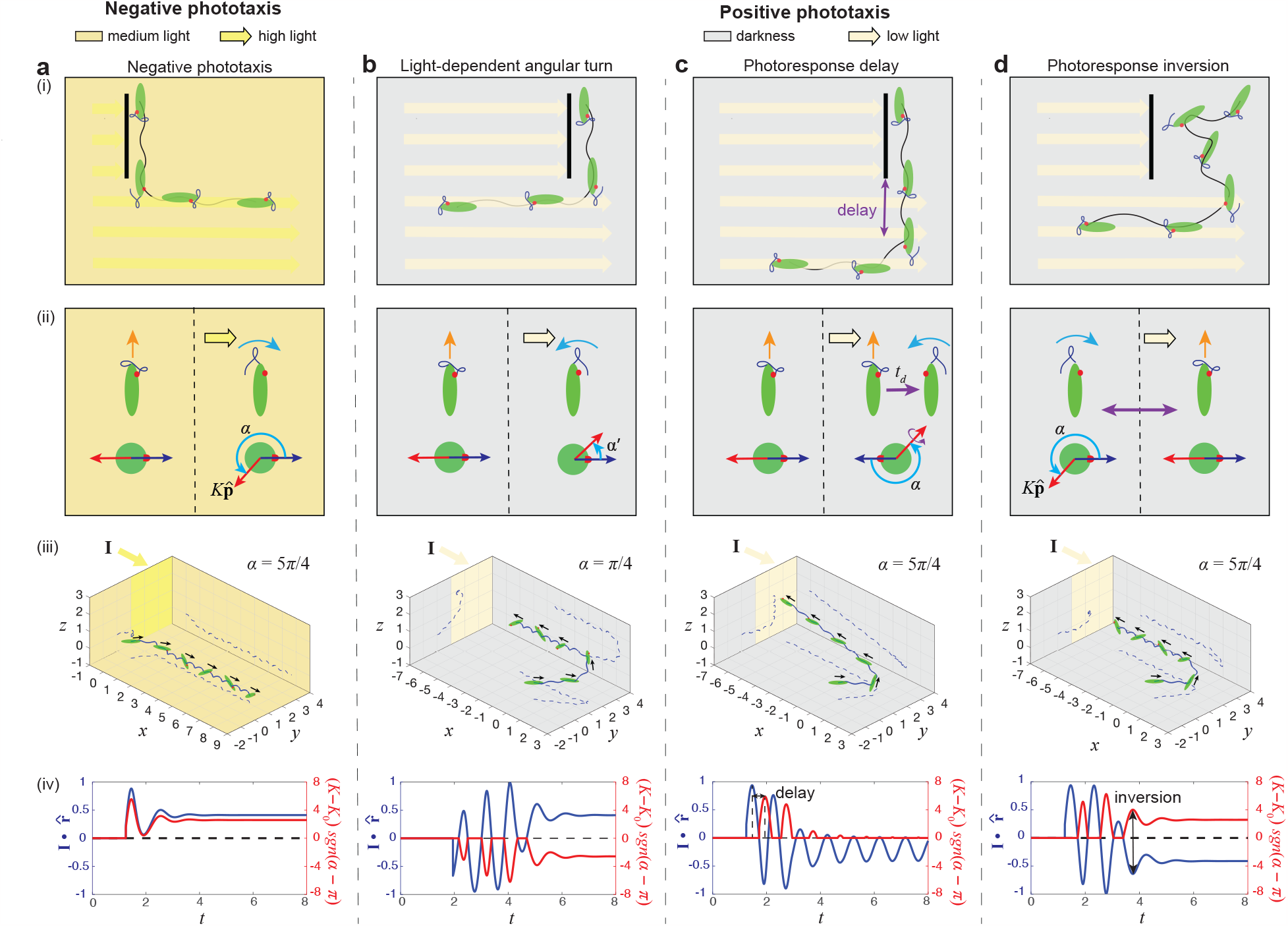
We propose and model three general mechanisms that could explain the transition between negative phototaxis and positive phototaxis. **(a)** Negative phototaxis is due to the tuning of paw angle *α* (Fig. 1e), where *E. gracilis* switches its beat pattern from forward swimming beats to sideward turning beats whenever it detects a strong light signal, leading to corresponding course corrections. **(b-d)** Positive phototaxis is hypothesized to be due to one of three possible general mechanisms: **(b)** Light-dependent angular turn suggests that *E. gracilis* tunes its paw angle *α* to a different direction as negative phototaxis and modulate its helical paths to achieve positive phototaxis. **(c)** Photoresponse delay suggests that *E. gracilis* achieves positive phototaxis by a delay mechanism in response to a light stimulus. **(d)** Photoresponse inversion suggests that *E. gracilis* inverts its response towards high light and low light, resulting in an opposite alternation of swimming and turning states between positive phototaxis and negative phototaxis. The panels in (i) illustrate typical swimming paths. The panels in (ii) illustrate the response of the flagellar beat patterns at different light intensities. The panels in (iii) show the simulation results. Initially, each cell swims into a directional light field **I** as indicated by the corresponding colored region in the z-plane; trajectories are projected onto the three planes with dashed lines. The panels in (iv) depict the light level as detected by the light sensor 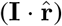as well as the turning response (*K −K*_0_)*sgn*(*α− π*) that represents the light dependent reorientation rate and thereby the direction of cell turning. See Supplementary Section 3.9 for details of model parameters).

To further substantiate the feasibility of these three possible mechanisms, we extended our previous biophysical model (Fig. 1e) [21] to also execute each of these mechanisms for positive phototaxis at low light intensities (Fig. 3b-d, Supplementary Sections 3.1 - 3.6). In our original model, the cell reorients with increased turning rate *K* (Fig. 1e) based on the detected light signal **I** · 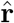 (light direction and intensity **I** and the photosensor pointing at 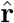). The model is non-dimensionalized in space and time based on the body length *𝓁* and the inverse of the rolling period 2*π/ω*, with *v* held constant. The coupling constants *K*_*d*_ are tuned such that the magnitude of *K*_*d*_|**I** |approximately equals to the light intensity in lux. If *α* is confined to *− ϵ/*2 *< α < ϵ/*2 and *π − ϵ/*2 *< α < π* + *ϵ/*2, with *ϵ≈* 0.2, then polygonal swimming behaviors in a plane orthogonal to the light direction are generated [21]. While not investigated explicitly in our previous publication [21], we now show that this model also already accounts for negative phototaxis for *π* + *ϵ/*2 *< α <* 2*π− ϵ/*2, with *ϵ≈* 0.2 (Fig. 3a(iii), Supplementary Movie S4), consistent with previous experimental results (Supplementary Sections 3.3 - 3.5) [11].

For the **light-dependent angular turn mechanism**, we adapted the model to modulate the turning vector 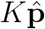 by making *α* dependent on light intensity. *K* depends on| **I**|and signal 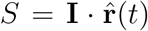 with corresponding coupling constants *K*_*a*_ and *K*_*d*_ for ambient and directional light components, respectively:

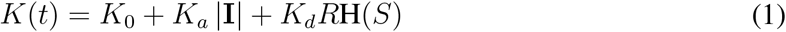

Here *K*_0_ is an intrinsic, light-independent reorientation rate and H is the Heaviside function accounting for eyespot shading. The response strength is given by *R* and is set to be equal to *S* here. The model exhibits positive phototaxis if *α* is confined to *ϵ/*2 *< α < π − ϵ/*2 (Fig. 3b(iii), Supplementary Movie S5), again with *ϵ ≈* 0.2.

For the **photoresponse delay mechanism**, we included a delay in signal transmission into the model, which is described by replacing the signal *S* with a delayed signal *S*_*d*_:

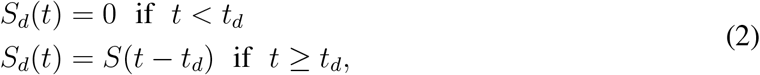

where *t*_*d*_ is the delay time of the response, and *t* = 0 is the time when the light first reaches the photoreceptor. For delays on the order of half a roll cycle the model then exhibits positive phototaxis (Fig. 3c(iii), Supplementary Movie S6).

For the **photoresponse inversion mechanism**, we introduced a phototaxis sign parameter *P* which captures this inversion of photoresponse due to light intensity increase vs. decrease over a threshold light intensity Δ:

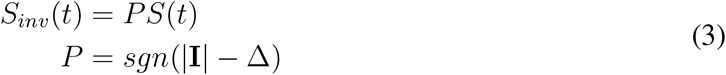

The cell inverts its photoresponse via a parameter *P*, where *P* =*−* 1 for positive phototaxis and *P* = 1 for negative phototaxis. The cell reorients and exhibit positive phototaxis whenever the eyespot shades its photoreceptor and the cell detects darkness (i.e., 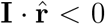, Fig. 3d(iii), Supplementary Movie S7).

Comparing the phase relation between *K* and 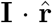 (for all three mechanisms blue curve in Fig. 3b-d(iv)), we find that for the first mechanism negative and positive phototaxis have the same phase relation (Fig. 3a(iv),b(iv), while the other two mechanisms show the opposite phase relation (Fig. 3c(iv),d(iv)). Importantly, for all three mechanisms the turning response (*K− K*_0_)*sgn*(*α−π*) is opposite to detected light 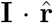 (red and blue curve in Fig. 3b-d(iv), respectively) in order to generate positive phototaxis (see Supplementary Section 3.4 for definition of turning response), while they are both aligned in order to generate negative phototaxis (Fig. 3a(iv)). In the following, we will experimentally test which of the three mechanisms is actually realized in *E. gracilis*.

### Experimental tests of the three mechanisms

To experimentally test the light-dependent angular turn mechanism (Fig. 3b), we measured the paw angle *α* for freely swimming cells under various conditions, i.e., (i) helical swimming in darkness (*∼* 0 lux), (ii) positive phototaxis under directional stimulus (⪅ 50 lux), (iii) helical swimming under medium background (*∼*100 lux), (iv) negative phototaxis under high stimulus (*>* 2, 000 lux) (Fig. 4a-d, Supplementary Section 4.3). In the two control cases of helical swimming under darkness (i) and medium background light (iii), *α* lies around *π* (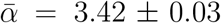 and 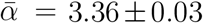), respectively, which agrees with the model predictions as *α* = *π* gives no bias towards or away from light. For negative phototaxis (iv), we obtained 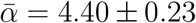, consistent with the expected range of *π* + *ϵ/*2 *< α <* 2*π ϵ/*2. However, for positive phototaxis (ii), we obtained (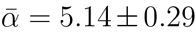), which is not significantly difference from *α* measured from negative phototaxis (*p <* 0.0001), and does not agree with the expected range of *ϵ/*2 *< α < π ϵ/*2 (*p <* 0.0001). This rules out the light-dependent angular turn mechanism (Fig. 4e).

**Figure 4.**
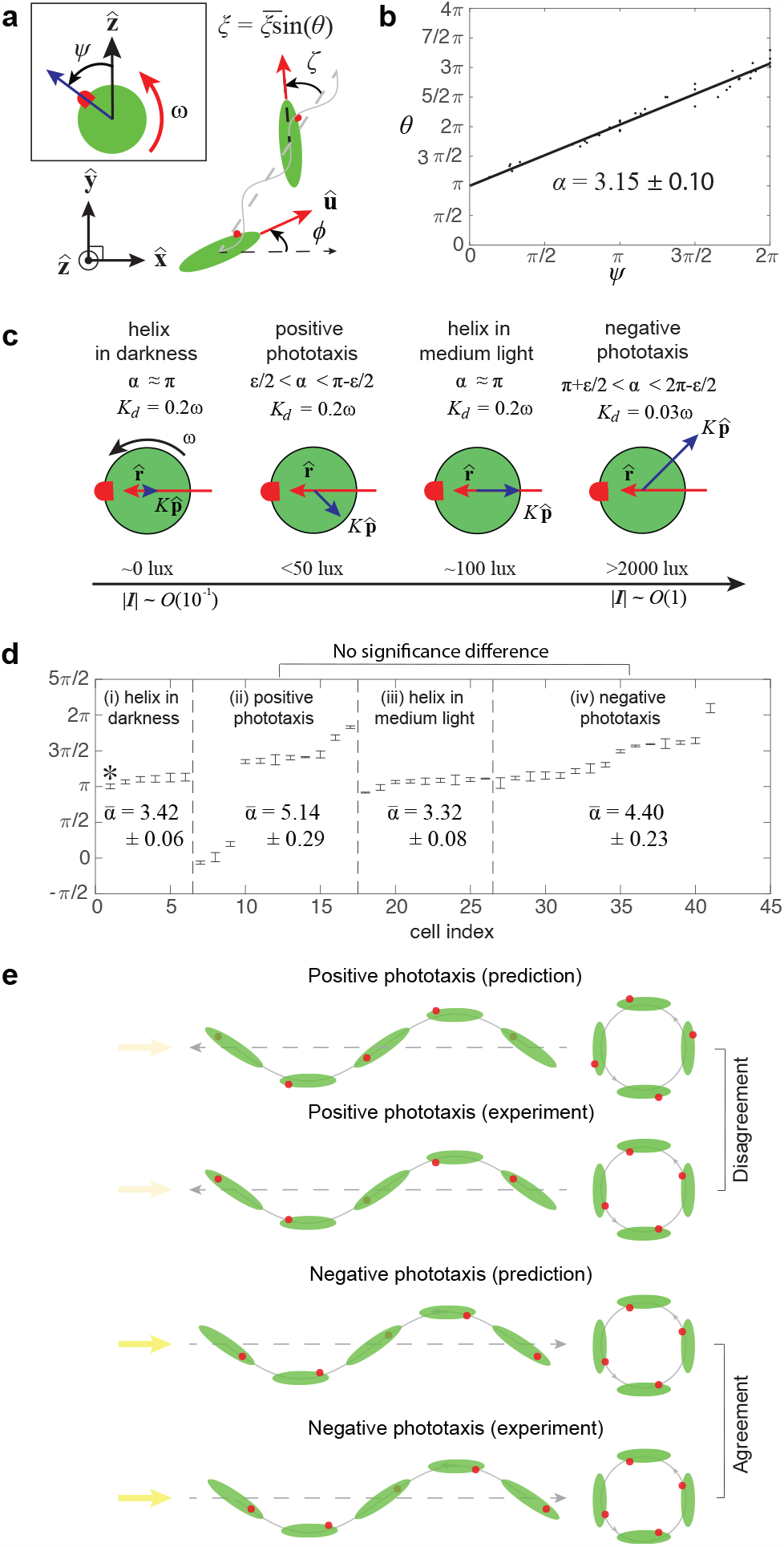
Experiments with freely swimming cells reveals that the Light dependent angular turn mechanism is not valid for *E. gracilis*. **(a-d)** Tracking of freely swimming cells show that positive and negative phototaxis display a similar range of paw angle *α*. **(a)** Schematic defining *ψ* and *θ*. **(b)** Linear fit for the phase relation between *ψ* and *θ* gives *α* in (i) darkness, (ii) positive phototaxis, (iii) medium light and (iv) negative phototaxis. **(c)** A schematic depicts the tuning of *α* and 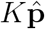 for different behaviors under the light-dependent angular turn mechanism. **(d)** *α* for different behaviors obtained from tracking of *N* = 41 cells for at least 3 rolling cycles each. The error bars denote the sem. **(e)** Schematics of predicted and experimentally observed eye spot orientation, which shows agreement for negative phototaxis but not for positive phototaxis.

To experimentally test the delay mechanism (Fig. 3c), we used micropipette aspiration [16] to fix *E. gracilis* cells in place (Fig. 5a) and measure the response of flagellar beat pattern upon changes in light intensity. To mimic conditions of positive and negative phototaxis, the same background and stimulus light conditions (i-iv) as in Fig. 4d were used, except for (iv) the stimulus was *>* 5, 000 lux (to make sure no cells falling into transitional polygonal behavior at intermediate light intensities [21]). Negative or positive phototaxis is triggered by tuning the microscope light intensity directly or applying an LED stimulus under a red observing light, respectively. We measured the time delay between the light stimulus turning on and the first beating state switching afterwards (Fig. 5d). We found for the negative phototaxis condition a delay of *t*_*d*_ = 0.23*±*0.06 s, while for the positive phototaxis condition the delay of *t*_*d*_ = 24.06 6.21 s was significantly larger (*n* = 15, *p <* 0.0001). We also observed a very wide distribution of delays ranging from a minimum of 0.24 s to a maximum of 55 s for the positive phototaxis condition. Importantly, we did not find a consistent delay difference between the positive and negative phototaxis condition on the time scale of half a roll cycle, i.e., *∼* 0.5 s, as would be required for inverting the swimming direction according to the delay mechanism (Fig. 3c). These results rule out the delay in photoresponse mechanism (Fig. 5e,f).

**Figure 5.**
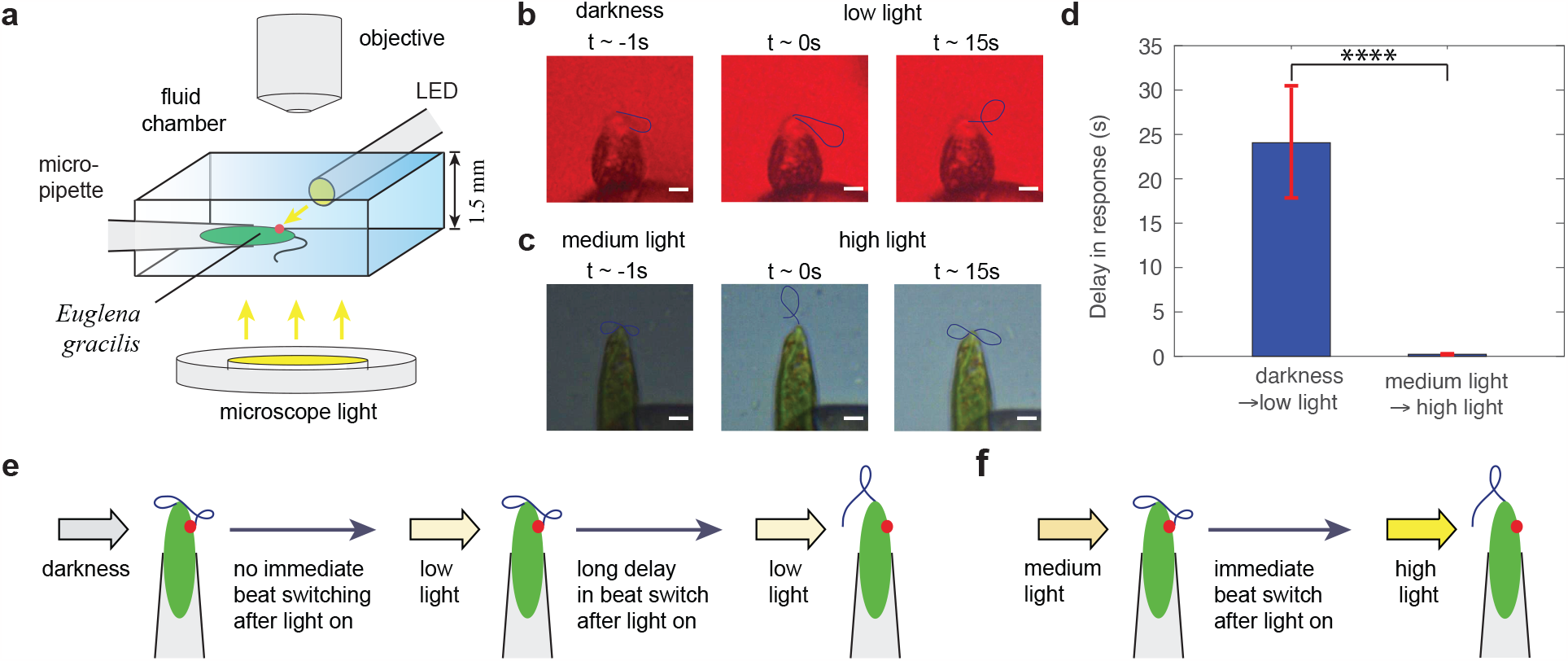
Experiments with spatially fixed cells reveals that the Photoresponse delay mechanism is not valid for *E. gracilis*. **(a)** Schematic of the experimental setup for a *E. gracilis* held on a micropipette. **(b)** Procedure for measuring time delay for switching the beat state from darkness to low light intensities. **(c)** Procedure for measuring time delay for switching the beat state from medium light to high light intensities. **(d)** Delay in response during light step-up. The errorbars represent the s.e.m. for the *n* = 15 samples from *N* = 6 cells. **(e)** Schematics of the switching in beat state and the corresponding time delay observed from darkness to low light. **(f)** Schematics of the switching in beat state and the corresponding time delay observed from medium light to high light. Scale bars in b,c: 5 *µ*m.

To experimentally test the photoresponse inversion mechanism (Fig. 3d), the observed beat patterns were quantified with the angle *λ* between the cell tip and the farthest material point on the flagellum within a beat cycle (Fig. 6a). We first measured the probability of the beating state switching of cells held in place upon changes in light intensity (Fig. 6b), which captures the proportion of turning beat patterns within 5 s after light changes. For the the negative phototaxis condition we found that the cells have a significantly higher probability of beat switching due to a step-up in light intensity (*n* = 15, *p <* 0.0001), while for positive phototaxis condition, the cells have a significantly higher probability of beat switching due to a step-down in light intensity (*p <* 0.0001). We then investigated the transient and adaptation behavior by periodically turning the stimulus on and off, with the on-period and the off-period lasting at least half a minute (Supplementary Section 4.4). In all cases, the cells displayed the two main beat patterns corresponding to helical swimming and turning behavior (Figs. 1d, 6a) as also established previously [21]. Under negative phototaxis light condition, the flagellum exhibits a swimming beat pattern under medium background light, while switching to the turning beat patterns almost immediately after exposure to a high stimulus light (Fig. 6c(i), Supplementary Movie S8). When turning the stimulus light off, the beat state switched back to the swimming state (Fig. 6c(ii), Supplementary Movie S9). Under the positive phototaxis condition, the flagellum stays at a swimming state when the LED is turned on, but once the LED was turned off the flagellum switched between the two beat patterns stochastically (Fig. 6c(iii), (iv), Supplementary Movie S10, Supplementary Movie S11). Hence the responses to changes in light intensity are fundamentally inverted between the negative and positive phototaxis conditions, i.e., the response is most robust and predictable as well as most immediate when the light intensity is increased vs. decreased, respectively (Fig. 6b). Furthermore, the cell responds with the turning vs. swimming beating pattern, respectively, once it perceives an increased light stimulus (Fig. 6c(i) and Fig. 6c(iii)). We also observed stochastic switching and adaptation of beat patterns when the light intensity was held constant over longer times after an initial change in light intensity (Supplementary Section 4.4). All experimental results together are consistent with the prediction of the photoresponse inversion mechanism (Fig. 3d).

**Figure 6.**
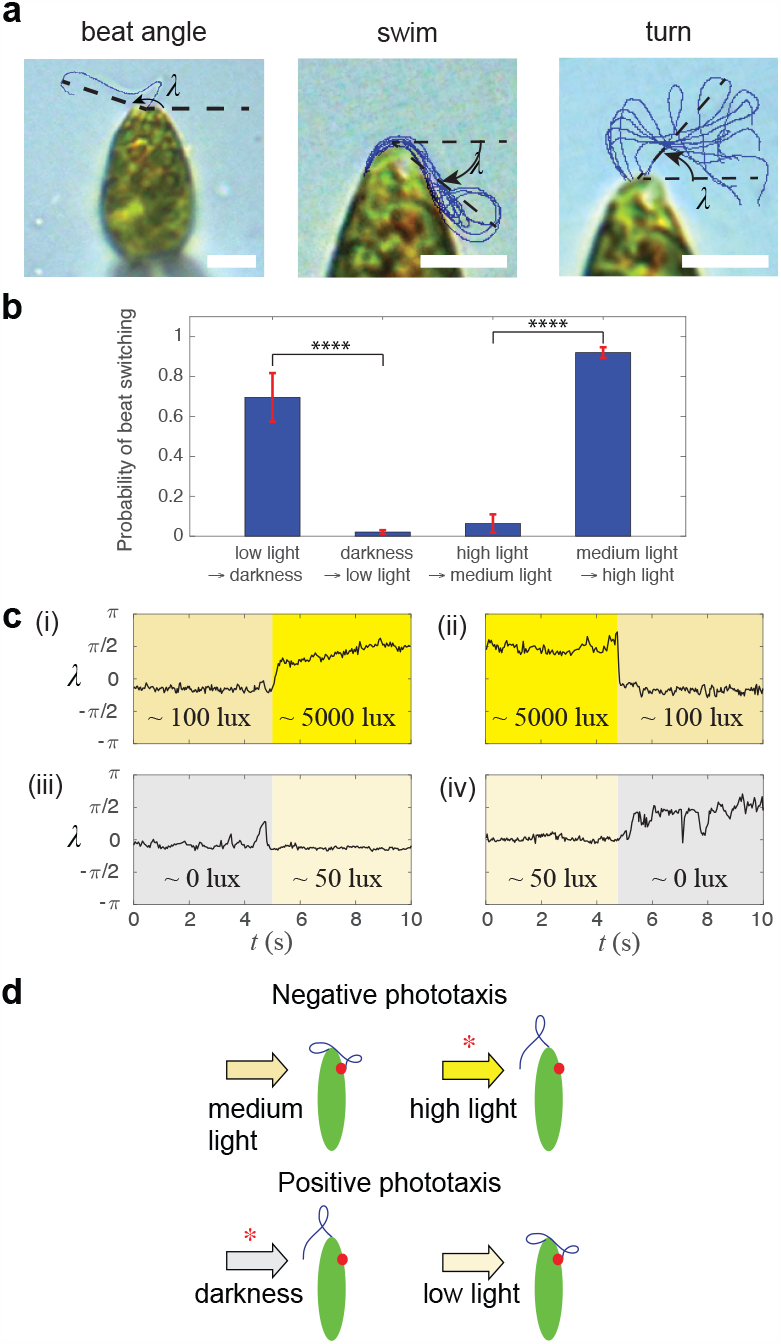
Experiments with spatially fixed cells reveals that the photoresponse inversion mechanism is consistent with *E. gracilis* light responses. **(a)** A beat angle *λ* is defined to quantify flagellar beats (left). Two main beat patterns corresponding to swimming (middle) and turning (right) were observed, and that are equivalent to those observed previously (Fig. 1d, [21]). Scale bars: 5 *µ*m. **(b)** Probability of cells switching their beat patterns after light step-up and light step-down. The errorbars represent the s.e.m. for the *n* = 15 samples from *N* = 5 cells. **(c)** Measurement of *λ* before and after light step-up and light step-down between different light intensities: 0 lux (grey), 50 lux (blue), 100 lux (pale yellow), 5, 000 lux (yellow); typical traces shown. **(d)** *E. gracilis* coordinates between two beat patterns at different light intensities to achieve positive and negative phototaxis strategies, i.e., *E. gracilis* utilizes the ‘photoresponse inversion mechanism’ (Fig. 3d); the red asterisks indicate an ‘activated turning state’ due to changes in light intensity.

In conclusion, among the three initially hypothesized mechanisms (Fig. 3d) the inversion mechanism is experimentally well supported for *E. gracilis*, while the other two mechanisms are ruled out for this organism. This also highlights an interesting solution for light searching based on a two-state feedback dynamics that applies to both positive and negative phototaxis under very high and very low light conditions, respectively (Fig. 6d): The cell mainly swims with helical beats in its favorable light conditions (low or medium background light), and switches to turning beat patterns for reorientation when it experiences unfavorable light conditions (darkness or high light). This simple feedback mechanism then enables effective phototaxis strategies over a very large range of light intensities (i.e., from 0 lux to over 10, 000 lux).

### Theoretical performance of the three mechanisms

This raises the question of why the inversion mechanism is used in *E. gracilis*, and we hypothesize that it provides a significant performance advantage under relevant conditions (and potentially it might also be easier to implement on a molecular or cellular level, and it might be easier to evolve). To test that, we quantified the experimentally measured mean free path and reorientation rate of the biased random walk of *E. gracilis* during positive phototaxis: Compared to the cells in darkness, the positively phototactic cells have a shorter mean free path, indicated by the rapid decrease in correlation of displacement direction (i.e., smaller *C*_*τ*_ over times, *n* = 50, *p <* 0.01, Fig. 7a, Supplementary Section 4.2). Moreover, the positively phototactic cells reorient more frequently, as indicated by the wider distribution of the reorientation rate (i.e., larger variance, *n* = 50, *p <* 0.0001, Fig. 7b, Supplementary Section 4.2). Hence noise seems to play a significant role during positive phototaxis.

**Figure 7.**
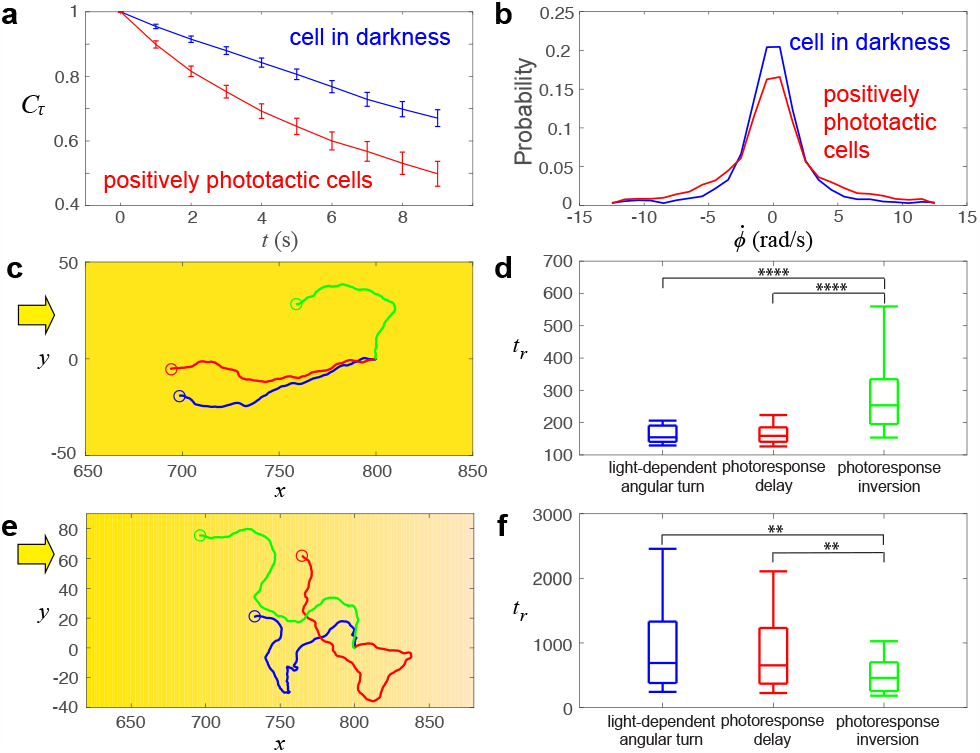
Depending on environmental and stimulus conditions, the photoresponse inversion mechanism has superior navigational performance than the other two mechanism as it can better overcome the sensory noise from its photoreceptor for positive and negative phototaxis. **(a)** Experimental measurement demonstrates that *E. gracilis* cells exhibiting positive phototaxis display a smaller displacement direction correlation *C*_*τ*_ over times (Supplementary Section 4.2) and **(b)** have a wider distribution of cell reorientation rate compared to cells in darkness. **(c)** Under parallel light rays, cells following the light-dependent angular turn (blue color) and photoresponse delay (red color) reach a light source faster than a cell following the photoresponse inversion mechanism (green color). **(d)** The box plots of *t*_*r*_ for the three mechanisms for the case of parallel light rays shown in (c). **(e)** Under a weak light gradient, simulations show that a cell following the photoresponse inversion mechanism (green color) typically outperforms the cells following the light-dependent angular turn (blue color) and photoresponse delay (red color) to reach a light source. **(f)** The box plots of *t*_*r*_ for the three mechanisms for the case of a weak light gradients as shown in (e). In (d) and (f), the midlines represent the median, and the box’s upper and lower bounds indicate the interquartile range; the upper and lower whiskers denote the 9th and 91st percentile of the simulation data, respectively. 100 simulations were performed for each mechanism.

We then considered the typical noise types and levels involved in positive phototaxis and the resulting effects on the swimming paths and ultimately on taxis behavior. The light intensity during positive phototaxis in our experiments (Fig. 2c,d) is *∼*50 lux and thereby potentially reflects natural settings, and which corresponds to 2*×*10^6^ photons/s hitting the photoreceptor (Supplementary Section 3.7). There is evidence that flavoproteins and flavins are responsible for *E. gracilis*’s photoreception [11], which feature a low absorption rate of photons (*∼*1%) [49], thus the effective light signal would be 2*×*10^4^ photons/s. Moreover, the thermal noise for a flavin-based photoreceptor is around 10^4^ photons/s [50], which is comparable to the effectively detected light signal. In contrast, the light signal for negative phototaxis (*>* 2, 000 lux [9, 23, 24]) is typically 80 fold larger compared to the sensory noise, which is then likely sufficient for the cell to discriminate the signal direction from noise and therefore exhibit directional response (Fig. 2a,b). We also estimate that the Brownian noise does not play a significant role given the comparably large cell body size of *E. gracilis*, e.g., the rotational diffusion due to thermal noise is *D*_*r*_ *∼*1.5*×*10^*−*6^ rad/s (Supplementary Section 3.2), and even the rotational diffusion due to active reorientation of the cells in the dark was measured to be only 0.02 rad/s (Supplementary Section 3.2) while *E. gracilis* can make course corrections of about 0.35 rad within a single beat stroke, i.e., within 50 ms (see below). Hence we propose that the main noise limitation for *E. gracilis* cells during positive phototaxis at low light conditions stems from receptor noise, while for negative phototaxis at high light conditions noise does not appear to play any role; hence any search mechanism needs to particularly overcome these noise effects at low light conditions.

Based on these considerations, we incorporated translational and rotational diffusion as well as sensory noise into all three mechanisms and then simulated their phototaxis performance the under low light levels and low signal-to-noise levels (Supplementary Section 3.5 - 3.6):

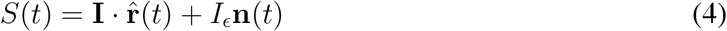

Here *S*(*t*) is the perturbed signal detected by the cell, *I*_*ϵ*_ is the noise strength and **n**(*t*) is a random vector with components following a Gaussian distribution with zero mean and unit variance. Based on the experiments shown in Fig. 6c and Supplementary Section 4.4, we now need to also account for the stochastic beat switching in the case of the photoresponse inversion model, hence we now use for *R* in Eq. (1):

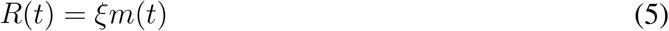

Here *ξ* sets the maximum response and *m*(*t*) is a random variable with a uniform distribution in [0, 1]. We then determined the theoretical performance of the different mechanisms by measuring the time *t*_*r*_ required for a cell to travel a specific distance of 100 cell body lengths towards the light source. We found that in the absence of attenuation of light, i.e., the light intensity |**I**| being constant everywhere, cells following the photoresponse inversion mechanism typically reach the goal slower than cells using the other two mechanisms (*n* = 100, *p <* 0.0001, Fig. 7c,d, Supplementary Movie S12). Conversely assuming that light intensity attenuates over ecologically relevant values (and as mimicked in our experimental conditions) (Supplementary Section 3.6), we found the photoresponse inversion mechanism performs significantly better than the other two (*n* = 100, *p <* 0.01, Fig. 7e,f, Supplementary Movie S13): The amount of light at a location **x** is given by combining Lambert’s law and inverse square law [51]:

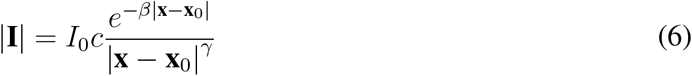

Here *I*_0_ is the light intensity at a reference location **x**_0_. *c, β* and *γ* are coefficients accounting for optical aberration, absorption and path loss. For typical ecological conditions as well as for our experimental conditions the parameters are *c* = 0.74, *β* = 0.29 and *γ* = 0.056, hence light intensity approximately decreases by 1*/*4 to 1*/*2 when doubling the distance (Supplementary Section 3.6).

From visually inspecting the simulated swimming trajectories, it appears that under a weak light condition the photoresponse inversion mechanism allows a cell currently swimming in a wrong direction to correct its swimming direction by a stochastic response, while the other two mechanisms heavily depend on detecting a sufficiently high light intensity and thereby fail to correct the swimming direction if the signal is too weak. These results suggest that the photoresponse inversion is a more robust mechanism under weak and noisy light conditions and an intrinsically noisy sensor.

## DISCUSSION

We proposed multiple theoretical mechanism of how microswimmers in principle could select between various taxis strategies based on light conditions (Fig. 3) [21], and we conclude that for *E. gracilis* the ‘photoresponse inversion’ is the only suitable one consistent with all data presented here and in the literature [10, 11, 24] (Fig. 8a). For high signal-to-noise levels as in the case of negative phototaxis, this mechanism leads to a directional steering mechanism by tuning the paw angle *α* (i.e., helical klinotaxis); while for low signal-to-noise levels as in the case of positive phototaxis, this leads to a run-and-tumble strategy (see further discussion below, [38])).

**Figure 8.**
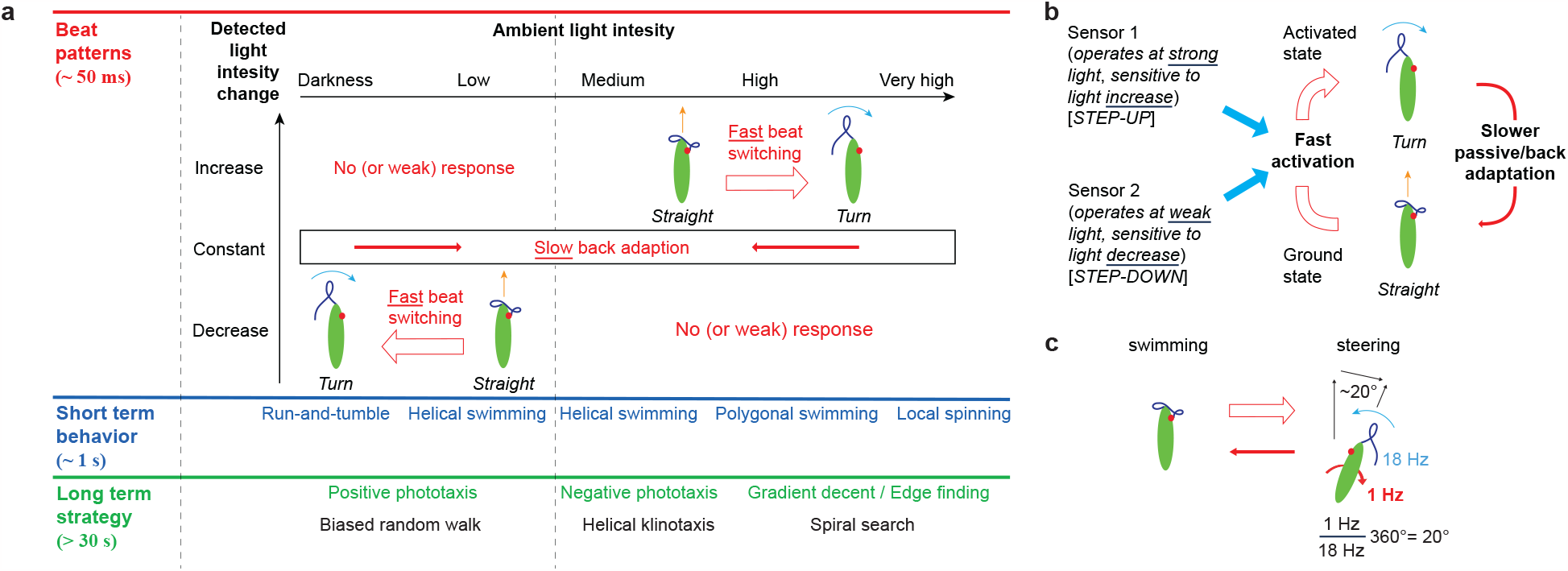
*E. gracilis* adapt the photoresponse inversion mechanism to overcome the sensory noise from its photoreceptor for positive and negative phototaxis. **(a)** Proposed behavioral phase space based on result of this paper and the much wider previous literature. Switching between the two beating patterns at short time scales leads to various medium term photobeaviors and various long term phototaxis strategies. **(b)** ‘Two sensor / two-beat state (inversion) model’: Two photosensors actuate the ‘turning beating state’, where one actuates based on light decrease at very low intensity, and the other actuates on light increase at high light intensity. The switching back to the ‘forward swimming beating state’ likely happens passively on slower times scale by adapting to the changed light intensity over time (here we hypothesize that the back-switching cannot be directly actuated, but a reverting back to the original light intensity might speed up the adaptation process). **(c)** Illustration on how discreteness of the beat as well as the amount of angular change per beat cycle relative to the number of beats for one full roll ultimately determines the orientational steering resolution (as also had already been found experimentally by others).

Importantly, the underlying flagella responses at low vs. at high light intensities are inverted, i.e., at low intensity flagella beating primarily reacts to a decrease in light intensity (’step-down’) vs. at high intensity to an increase in light intensity (’step-up’) [11, 22, 37]. In both cases, the ‘step-response’ represents a fast actuation from a ‘forward swimming beat state’ (’ground state’) to a ‘turning beat state’ (’activated state’), and with a (likely passive) back-relaxation over a typically longer time scale (Fig. 8b). The actuation and relaxation time scales might be different for both step-responses as different photorecptors are likely involved [52, 53], but we determined for the step-up responses 0.23*±*0.06 s (consistent with [40]) and 25.3*±*6.53 s, and for the step-down responses 1.02*±*0.26 s and 24.07*±*6.21 s. Many of the earlier reports relied on whole cell observations that did not resolve the flagallar beating states given understable limitations in imaging technology at the time [11, 37], but which we resolved now ([21] and Figs. 5b, 6a). Hence our work now closes this gap in understanding between step responses and flagellar beat patterns, with key implications for the resulting steering and search strategies (Fig. 8a).

How the selection of these different photobehaviors at different light intensity scenarios is achieved at a sub-cellular and molecular signaling cascade level is not fully understood and beyond this work [32, 33, 52–54]. But in short, there is strong evidence that positive and negative taxis are regulated by different flavin-based photosensors, e.g., a family of dissimilar Photoactivated adenylyl cyclases (PACs), and that they then likely feed into the same flagella activation mechanism [32, 33, 52–54]. Having two different photosensors might be advantageous for at least two reasons: (1) being optimally responsive to a decrease vs. an increase in light intensity (i.e., step-down vs. step-up, respectively), and (2) being optimally sensitive to very low vs. very high light intensities.

What then determines (and limits) the taxis performance and steering precision (Figs. 2b,c; 7b)? We see multiple factors, i.e., *E. gracilis* operates with discrete beat strokes leading to discrete (’quantized’) steering, the cellular motion is subjected to translational and rotational Brownian motion, the photosensors and molecular signaling cascades are subjected to a certain noise level including photon-shot-noise, and the light stimulus might be noisy.

Regarding the performance limits for negative phototaxis under high light intensity, it was reported previously that *E. gracilis* can correct its path by*∼*20^*°*^/s [11]. We note that this resolution limit can be directly understood from the flagellar beating characteristics (Fig. 8c): We previously reported that flagellar turning beats change the direction by*∼*18^*°*^ per beat [21]. At the same time, a cell executing negative phototaxis takes*∼*1 s to role around its long axis [21], and where only one of these beats per full role is used for a corrective turning beat while the photodector detects this corrective signal (assuming the cell is already in close alignment of the desired path), i.e., initiating a course correction of*∼*18^*°*^/s. Related arguments were summarized in [11], yet the connection apparently was not made that these*∼*20^*°*^/s correspond to a single flagellar beat. Yet, this course correction is experimentally harder to detect directly as it is embedded in the helical oscillatory motion. Our experiments showed that the distribution of reorientation rates is*∼*70^*°*^/s (see stdev in Fig. S8b), where cells with helical pitch of*∼*20^*°*^ will yield similar reorientation rates. We conclude, that at high light levels eventually lead to a primarily straight swimming path (with a superimposed helix) for negative phototaxis.

For performance limits for positive phototaxis, we pointed out above that the thermal noise for a flavin-based photoreceptor is around 10^4^ photons/s [50], which is comparable to the effectively detected light signal. For perfect steering we would again expect about*∼*20^*°*^/s (Fig. 8c), but given the sensor noise, a significant proportion of wrongly missed or added steering actions are expected, thereby limiting this resolution. By comparing the reorientation rates of cells in darkness (1 stdev = 1.88*±*0.14 rad/s, 95% C.I., Fig. 6b) with that of cells under positive phototaxis (1 stdev = 2.57*±*0.29 rad/s, 95% C.I., Fig. 6b), we can estimate the steering resolution to be*∼*40^*°*^/s, i.e., approximately two corrective beats per cycle. This result agrees with our estimation of steering resolution for positive phototaxis.

Note that with our estimates of performance limits, we consider corrective signal changes that are so low that the activation energy is very low and consequently the back-adaptation time is short, i.e., the ‘slow’ back adaption is essentially on the order of a single flagellar beat. Note that all our simulations so far had been based on a time-continuous response, i.e., no explicit model of discrete beat with discrete changes in cell orientation was assumed; we therefore then also implemented and tested such a discrete model, but all results and conclusions stayed the same (Supplementary Section 3.5). We conclude that for positive phototaxis *E. gracilis* operates at the noise limits of sensor detection (and potentially also internal signal transmission) leading to a swimming path representing a biased random walk [6].

Our results then also provide a rather concise explanation for the versatile and complex set of photo-behaviors of *E. gracilis* as described by us and the literature (Fig. 8a), such as positive and negative phototaxis along the light direction, polygonal search perpendicular to the light vector in complex light fields, transitioning between normal vs. anormalous diffusion, and localized spinning [9–11, 21, 22, 24, 24, 25, 30, 31, 37]. Our modeling shows how all these behaviors can be understood by switching between only these two beat states (Fig. 8a) in the right temporal sequence and based on the currently sensed light stimulus, and where adaptions in light sensing and/or signaling processing play an important role as well.

We also note that other photo-behaviors exist that are potentially not explained by just these two beating states, for example, localized spinning does not seem to involve any rotation around the long axis anymore as observed in all other beat states (although this could still be explained by a rotation around two axes where one is effectively canceled out [21]); there are also reports on backward swimming that likely require a substantial different beating pattern [55]; and other behaviors like photokinesis (i.e., the change in velocity based on light intensity changes[11]), swimming at an angle relative to the vector of polarized light [36, 56], or the swimming direction when subjected to multiple light sources from different directions [45]. Our model can already recapitulate some of these aspects (Supplementary Section 3.8), and all these behaviors deserve further experimental and theoretical study, which is beyond this paper and left for future work.

We also speculate that the biophysical constraints on phototaxis may explain the relative and absolute placement of eyespot and flagellum, and why *E. gracilis* is a puller-type swimmer [2]. Intuitively, it would be advantageous for the flagellum to be close to the photoreceptor to aid a fast and robust signaling. Furthermore, during negative phototaxis under highest light intensity the photosensory organ might be protected against photodamage (and also be more sensitive) by being shaded from the whole cell body, while during positive phototaxis the sensor points mostly towards the light source and would thereby be able to capture more of the scarce photons that would be adsorbed by the cell body otherwise. This design rule might be extensible to other phototactic cells such as Chlamydomonas [14, 16].

The three phototaxis mechanisms we proposed in this work (Fig. 3) may be generalizable and apply to other phototactic organisms. Microorganisms (e.g., Chlamydomonas and Gonium) or larger organisms (e.g., Platynereis) swim in helical paths during phototaxis to enhance their light sensing [12, 14, 57, 58]. We note that although we ruled out light-dependent angular turn for positive phototaxis of *E. gracilis*, the phase relationship captured by this mechanism seems to agree with both positive and negative phototaxis for Chlamydomonas reinhardtii, where the eyespot was reported to be located at the outer (inner) side of the helix during positive (negative) phototaxis, respectively [13]. We also expect that the delay in photoresponse mechanism may have implications for the selection of the phototaxis sign in other phototactic organisms [57]. A deeper biophysical comparison between these (and other) organisms would be valuable for future work.

Finally, our results can have profound implications not only for natural microswimemers, but also for the design and control of light-guided synthetic microswimmers [59]. The three proposed mechanisms (Fig. 3) may also provide effective solutions for developing synthetic swimmers [59– 62] that can select between positive and negative taxis in response to environmental stimuli (e.g., chemicals, light, flows, etc) using a simple noisy sensor and a two-state feedback response.

## Acknowledgements

We like to thank members of the Riedel-Kruse lab and R. Davis for useful discussions. This work was supported by NSF #1324753 (I.H.R-K), GRF #127208421 and Croucher Foundation (A.C.H.T.).

