## Supplementary Material for "Flagellar beat state switching in microswimmers to select between positive and negative phototaxis"

### Switching mechanisms between positive and negative phototaxis for chiral microswimmers

#### 1 Supplementary movies

**Movie S1:** Negative phototaxis experiment. The video is replayed at the real time.

**Movie S2:** Positive phototaxis experiment. The video is replayed at the real time.

**Movie S3:** Negative phototaxis experiment, individual cell shows beat switching. The images were sampled at 200 fps and the video is replayed at 10× slower than the real time.

**Movie S4:** Negative phototaxis simulation.

**Movie S5:** Positive phototaxis simulation, light-dependent angular turn.

**Movie S6:** Positive phototaxis simulation, delay in photoresponse.

**Movie S7:** Positive phototaxis simulation, inversion in photoresponse.

**Movie S8:** Micropipette experiment, medium light to strong light. The images were sampled at 200 fps and the video is replayed at 10× slower than real time.

**Movie S9:** Micropipette experiment, strong light to medium light. The images were sampled at 200 fps and the video is replayed at 10× slower than real time.

**Movie S10:** Micropipette experiment, dark to low light. The images were sampled at 200 fps and the video is replayed at 10× slower than real time.

**Movie S11:** Micropipette experiment, low light to dark. The images were sampled at 200 fps and the video is replayed at 10× slower than real time.

**Movie S12:** Simulations of the three mechanisms, parallel light.

**Movie S13:** Simulations of the three mechanisms, light gradient.

More details about the parameters used for the simulations in Movies S4-S7, S12, S13 can be found in section 3.9. For movies S4-S7 and movie S12-S13 (all simulations), the length scale and the time scale are taken as the body length and the roll period of the cell, respectively. The cell swims at a speed of one body length per roll cycle.

### 2 Supplementary methods

#### 2.1 Cell preparation

*Euglena gracilis* was obtained from Carolina Supplies (#152800). The cultures were kept in the original glass containers with the lids loosened or in an open reservoir at room temperature ($\sim$ 24°C) with a medium ambient light ($\sim$ 500 lx, recorded at region outside the glass containers). Light intensities were measured with a AEMC CA811 lightmeter. The illuminance (lux) was measured based on light stimulus illuminated on an area of $\sim$ 1.5 mm. The cell culture could be readily used for negative phototaxis experiments. For positive phototaxis experiments, the cells were kept in dark for 24 hours before the experiments such that the cells were fully adapted to darkness before any light stimuli was introduced. This preparation step enhanced the sensitivity of the cells towards weak light stimuli (see also section 4.1).

#### 2.2 Experimental setups

Details of the experimental setups are given by the followings:

**Flow chamber**

For experiments shown in main text Fig. 2 and Fig. 7a,b, the Euglena were situated in a $\sim$ 300 $\mu$m deep flow chamber formed by sandwiching two layers of double-sided tape strips between a glass slide and a cover slip. The two layers of strips was placed on each side of the flow chamber to set the effective height of the chamber measured with a caliber. The cells were imaged at 4$\times$ magnification for a wide field of view to capture the cell trajectories. For experiments shown in main text Fig. 4, the cells at 20$\times$ magnification inside a $\sim$ 300 $\mu$m deep microfluidic chamber were followed by moving the microscope stage by hand to track the phase relationship between eyespot and cell reorientation. For the micropipette aspiration experiments shown in main text Fig. 5 and Fig. 6, the cells were situated in a $\sim$ 1 mm deep flow chamber to allow enough space for moving the micropipette inside the chamber. The cells were captured by the micropipette controlled manually by a micromanipulator. The captured cells were imaged at 40$\times$ magnifications to observe the fine details of the flagellum.

**Light stimulation**

For all experiments, the cells were imaged using bright field microscopy on a Leica DM500 compound microscope. Light intensities were measured with a AEMC CA811 lightmeter at a location where the light entered the flow chamber.

For negative phototaxis experiments shown in main text Fig. 2c,d and Fig. 4, the cells were illuminated by a constant light field the microscope lamp to provide an ambient background light at medium light intensities (i.e., $\sim 100 - 300$ lux). A LED was used to provide a strong light stimuli (i.e., $\sim 2,000$ lux to $> 10,000$ lux) from one side of the flow chamber to trigger negative phototaxis of the cells.

For positive phototaxis experiments shown in main text Fig. 2a,b, Fig. 4 and Fig. 7a,b, the observing light from the microscope lamp was filtered in red by a 630 nm longpass glass filter purchased from Thorlabs, where Euglena cells have shown to be not responsive to the red filtered light [1]. A weak LED stimulus (i.e., $\sim$ 50 lux) was introduced from one side of the flow chamber to trigger positive phototaxis of the cells. All positive phototaxis experiments were performed in a dark room to minimize the effects of ambient light.

For the micropipette aspiration experiments shown in main text Fig. 5 and Fig. 6, different settings were used to provide four light conditions we considered, i.e., darkness ($\sim$ 0 lux), low light ($\sim$ 50 lux), medium light ($\sim$ 100 lux) and high light($\sim 5,000$ lux). The medium light and high light conditions were provided by finely tuning the microscope light directly. The dark condition was provided by using a red filtered observing light in a dark room. The low light condition was provided by a weak LED from one side of the flow chamber with a red observing light.

**Imaging**

A Pixelink PL-D725CU-T camera was used for recording cell motions, where different imaging rates were used for different experiments. For experiments shown in main text Fig. 2 and Fig. 7a,b where we investigated the statistics of cell trajectories, cell motions were recorded at imaging rate of 10 fps for long times (up to 10 minutes). For experiments shown in main text Fig. 4 where the eyespot and orientation of swimming cells were tracked, cell motions were recorded at imaging rate of 30 fps for a typical record time of 30 s to 1 minute (until the track of the cells were lost). For micropipette aspiration experiments shown in main text Fig. 5 and Fig. 6, flagella motions of fixed cells were recorded at high imaging rate of 200 fps or 400 fps for 1 to 2 minutes.

### 3 Biophysical model

#### 3.1 Numerical implementation

For completeness, here the details of the numerical implementation of the model are included, and which can also be found in the supplementary materials of our previous publication [2].

The cell body rotation was modeled through quaternion rotation. For rotation about a unit vector $\hat{\mathbf{w}} = (w_x, w_y, w_z)$ with an angle $\Phi$, the quaternion is given by

$$\begin{aligned} q(\hat{\mathbf{w}}, \Phi) &= [q_0, q_1, q_2, q_3] \\ &= [\cos(\Phi/2), \sin(\Phi/2)w_x, \sin(\Phi/2)w_y, \sin(\Phi/2)w_z], \end{aligned} \tag{S1}$$

and the corresponding rotation matrix is

$$\mathbf{R} = \begin{bmatrix} 1 - 2q_2^2 - 2q_3^2 & 2(q_1q_2 - q_0q_3) & 2(q_1q_3 + q_0q_2) \\ 2(q_1q_2 + q_0q_3) & 1 - 2q_1^2 - 2q_3^2 & 2(q_2q_3 - q_0q_1) \\ 2(q_1q_3 - q_0q_2) & 2(q_2q_3 + q_0q_1) & 1 - 2q_1^2 - 2q_2^2 \end{bmatrix}. \tag{S2}$$

The cell rotation in a single time step $\delta t$ can be decomposed into a sequence of quaternion rotations: (1) reorientation due to light stimulus at yitch rate $K$; (2) cell rolling about its long axis at frequency $\omega$; and (3) reorientation due to rotational diffusion with a coefficient $D_r$ and a noise parameter $\xi_r$ (chosen from a normal distribution with zero mean and unit variance) about a uniformly random unit vector $\hat{\mathbf{f}}$ spanning over the 3D orientation space. To characterize the body rotation, the rotations of the long body axis $\hat{\mathbf{u}}$, direction of the maximum light sensitivity $\hat{\mathbf{r}}$, and the paw axis $\hat{\mathbf{p}}$ are required. The rotation sequence can be expressed as:

$$\begin{aligned} \hat{\mathbf{u}}_1 &= \mathbf{R}(\hat{\mathbf{p}}, K\delta t)\hat{\mathbf{u}}, \\ \hat{\mathbf{r}}_1 &= \mathbf{R}(\hat{\mathbf{p}}, K\delta t)\hat{\mathbf{r}}, \\ \hat{\mathbf{r}}_2 &= \mathbf{R}(\hat{\mathbf{u}}_1, -\omega\delta t)\hat{\mathbf{r}}_1, \\ \hat{\mathbf{p}}_1 &= \mathbf{R}(\hat{\mathbf{u}}_1, -\omega\delta t)\hat{\mathbf{p}}, \\ \hat{\mathbf{u}}(t+\delta t) &= \hat{\mathbf{u}}_1 + \mathbf{R}(\hat{\mathbf{f}}, \sqrt{2D_r\delta t}\xi_r)\hat{\mathbf{u}}_1 \\ \hat{\mathbf{r}}(t+\delta t) &= \hat{\mathbf{r}}_2 + \mathbf{R}(\hat{\mathbf{f}}, \sqrt{2D_r\delta t}\xi_r)\hat{\mathbf{r}}_2 \\ \hat{\mathbf{p}}(t+\delta t) &= \hat{\mathbf{p}}_1 + \mathbf{R}(\hat{\mathbf{f}}, \sqrt{2D_r\delta t}\xi_r)\hat{\mathbf{p}}_1 \end{aligned} \tag{S3}$$

Defining the center of the cell as $\mathbf{c}$ and its swimming speed as $v$, the translational motion of the cell is given by

$$\mathbf{c}(t+\delta t) = \mathbf{c}(t) + (v\hat{\mathbf{u}}(t))\delta t + \sqrt{2D_t\delta t}\xi_t\hat{\mathbf{g}}. \tag{S4}$$

where $D_t$ is a translational diffusion coefficient and $\xi_t$ is chosen from a normal distribution with zero mean and unit variance. $\hat{\mathbf{g}}$ is a uniformly random unit vector spanning over the 3D orientation space.

Given the response function that accounts for the change in $K$ over time (main text Eqs. (1)–(6)), together Eqs. (S1)–(S4) form a dynamical system that describe the control feedback between eyespot, cell orientation, light detection, light adaptation and cellular reorientation, effectively capturing the phototaxis of Euglena.

##### 3.2 Translational and rotational diffusion noises

Here a summary is provided on the various sources of noises for translational and rotational diffusion of Euglena, and their corresponding magnitudes are compared.

First the noises due to thermal diffusion is considered. If the anisotropy of the cell body is neglected and the effective diffusion due to a characteristic length $\ell$ of the cell (i.e., $\ell \sim 50\mu$m, typical length of a Euglena cell) is considered, the translational and rotational noises due to the thermal diffusion can be captured by the Stokes-Einstein relation [3]:

$$\begin{aligned} D_{t,thermal} &= \frac{k_B T}{6\pi\eta\ell}, \\ D_{r,thermal} &= \frac{k_B T}{8\pi\eta\ell^3}, \end{aligned} \tag{S5}$$

where $k_B = 1.380649 \times 10^{-23}$ is the Boltzmann constant, $T$ is the reference temperature in Kelvin scale and $\eta$ is the dynamic viscosity of water. At a room temperature of $T = 298$ K and $\eta = 8.9 \times 10^{-4}$ kg/m·s, $D_{t,thermal} \sim 2 \times 10^{-6}$ $\mu$m$^2$/s and $D_{r,thermal} \sim 1.5 \times 10^{-6}$ rad/s. Noises due to $D_{t,thermal}$ and $D_{r,thermal}$ are negligible compared to the velocity $v$ of Euglena (i.e., $v \sim 50$ $\mu$m/s) and the effective diffusion $D_{eff}$ of non-stimulated Euglena cell measured in previous experiments (i.e., $D_{eff} \sim 2 \times 10^4$ $\mu$m$^2$/s) [2]. The effective diffusion is hence due to active reorientation of the cell and flagellar noise.

The relationship between the effective diffusion $D_{eff}$ and the rotational noise due to active reorientation of the cell $D_{r,active}$ is given by the Taylor dispersion theory [4]:

$$D_{eff} = \frac{v^2}{6D_{r,active}}. \tag{S6}$$

Given $D_{eff} \sim 2 \times 10^4$ $\mu$m$^2$/s and $v \sim 50$ $\mu$m/s for Euglena, $D_{r,active} \sim 0.02$ rad/s. The translational and rotational noises are considered in Eq. S3 and Eq. S4 via $D_t = D_{t,thermal}$ and $D_r = D_{r,thermal} + D_{r,active}$.

##### 3.3 General model of photoresponse

Here a general model of photoresponse is considered that accounts for sensory noise and response strength. In our model, the Euglena cell rotates along a "yitch-paw" axis in response to light stimuli (main text Fig. 1e). The strength of the "yitch-paw" axis is denoted by a yitch rate $K$ and the angle between the eyespot vector and the "yitch-paw" axis is denoted by a paw angle $\alpha$.

The yitch rate $K$ depends on depends on $|I|$ and and signal $S$ $(\mathbf{I} \cdot \hat{\mathbf{r}}(t))$ with corresponding coupling constants $K_a$ and $K_d$ for ambient and directional light components, respectively:

$$K(t) = K_0 + K_a\,|\mathbf{I}| + K_d R \mathrm{H}(S) \tag{S7}$$

Here $K_0$ is an intrinsic, light-independent reorientation rate and H is the Heaviside function accounting for eyespot shading. $S$ is the signal and $R$ is the response strength.

The signal $S$ depends on the detected light $(\mathbf{I}\cdot\hat{\mathbf{r}})$and the sensory noise by

$$S(t) = \mathbf{I}\cdot\hat{\mathbf{r}}(t) + I_\epsilon \mathbf{n}(t) \tag{S8}$$

Here $I_\epsilon$ is the sensory noise strength and $\mathbf{n}(t)$ is a random vector with components following a Gaussian distribution with zero mean and unit variance.

Depending on how the signal $S$ and the response strength $R$ are treated, there are three possible mechanisms for positive phototaxis in this model:

(1) Light-dependent angular turn: The cell turns by rotating along the "pitch-yaw" axis and the phototaxis sign is fully determined by the paw angle $\alpha$.

$S$ follows Eq. S8 and $R$ is light-dependent, i.e., $R = S$. Positive phototaxis occurs when $\epsilon/2 < \alpha < \pi - \epsilon/2$, negative phototaxis occurs when $\pi + \epsilon/2 < \alpha < 2\pi - \epsilon/2$. If $\alpha$ is confined to $-\epsilon/2 < \alpha < \epsilon/2$ and $\pi - \epsilon/2 < \alpha < \pi + \epsilon/2$, with $\epsilon \approx 0.2$, then polygonal swimming behaviors in a plane orthogonal to the light direction are generated [2].

(2) Photoresponse delay: There is a delay in signal transmission, which is described by replacing the signal $S$ in Eq. S8 with a delayed signal $S_d$:

$$\begin{aligned} S_d(t) &= 0 \ \text{ if } \ t < t_d \\ S_d(t) &= S(t - t_d) \ \text{ if } \ t >= t_d, \end{aligned} \tag{S9}$$

where $t_d$ is the delay time of the response. The response strength is given by $R = S_d$. $\alpha$ follows the range $\pi + \epsilon/2 < \alpha < 2\pi - \epsilon/2$.

(3) Photoresponse inversion: A phototaxis sign parameter P is introduced to capture the inversion of up-down photoresponse over a threshold light intensity $\Delta$:

$$\begin{aligned} S_{inv}(t) &= PS(t) \\ P &= sgn(|\mathbf{I}| - \Delta) \end{aligned} \tag{S10}$$

The cell inverts its photoresponse via a parameter $P$, where $P = -1$ for positive phototaxis and $P = 1$ for negative phototaxis. $\alpha$ follows the range $\pi + \epsilon/2 < \alpha < 2\pi - \epsilon/2$. The response strength is stochastic, given by

$$R(t) = \xi m(t) \tag{S11}$$

Here $\xi$ sets the maximum response and $m(t)$ is a random variable with a uniform distribution in $[0, 1]$.

#### 3.4 Turning response

In the absence of noises, the equilibrium angle $A_e$ where the cell swim with respect to light after reorientation is dependent on the choice of $\alpha$. An equilibrium angle is defined as the angle between by the light stimulus $\mathbf{I}$ and the average displacement vector of the cell trajectory after the reorientation, given by $\hat{\mathbf{x}}_{t\to\infty} = \frac{\int_{T_n}^{T_n+1}\mathbf{x_n(t)}dt - \int_{T_n}^{T_n-1}\mathbf{x_n(t)}dt}{|\int_{T_n}^{T_n+1}\mathbf{x_n(t)}dt - \int_{T_n}^{T_n-1}\mathbf{x_n(t)}dt|}$, where $T_n - T_{n-1}$ represents the duration of the $n$-th rolling period. Hence $A_e = \cos^{-1}(\mathbf{I}\cdot\hat{\mathbf{x}}_{t\to\infty}/|\mathbf{I}|)$. Fig. S1 shows $A_e$ for the three phototaxis mechanisms, where the regions of $\pi/2 < A_e < \pi$ corresponds to positive phototaxis and $0 < A_e < \pi/2$ corresponds to negative phototaxis. It can be seen that for all three models, the phototaxis sign switches when $\alpha$ shifts by $\pi$. Therefore, it is convenient to introduce a turning response $(K - K_0)sgn(\alpha - \pi)$ to compare the phototactic turning along the yitch-paw axis during positive and negative phototaxis. The first component $K - K_0$ will determine the turning rate with respect to the light-independent reorientation rate $K_0$. The second component $sgn(\alpha - \pi)$ determines the direction of turning with respect to the eyespot vector $\hat{\mathbf{r}}$, where the direction of turning changes to the opposite sign when shifting $\alpha$ by $\pi$. Therefore, this turning response account for both the strength and direction of the photoresponse in different model. The comparison of the turning response for different models are presented in Fig. 3a-d(iv) in the main text.

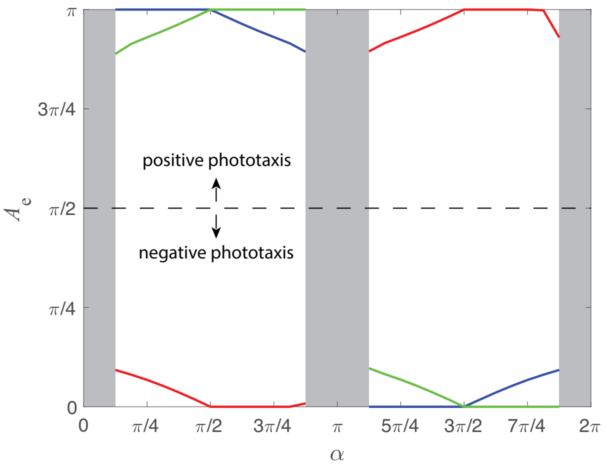


Figure S1: **Equilibrium angle $A_e$ of phototaxis given by different models with different paw angle** $\alpha$. The blue, red and green lines correspond to the phototaxis mechanism of light-dependent angular turn, photoresponse delay and photoresponse inversion, respectively. The shaded regions represent the transition case where the cell swims in a polygonal trajectory orthogonal to the light. Light stimulus is given by $\mathbf{I} = \hat{\mathbf{x}}$.

#### 3.5 Discrete vs. continuous response

Here the influence of implementing discrete response in our model is discussed. In general, Euglena adjusts its swimming direction discretely by a flagella beat. Therefore, a discrete photoresponse is perhaps a more general model to describe the cell dynamics due to discrete flagellar beats compared to a fully continuous model.

A discrete model is considered with 20 photoresponses within 1 roll cycle of Euglena (i.e., flagella beat frequency $\sim 20$ Hz for a cell with rolling frequency of $\sim 1$ Hz). The cell detects an instantaneous light stimulus at time $t$ and uses such stimulus as the signal for a discrete response. For the n-th discrete response, the signal is given by

$$S(t) = S(t_n) \quad \text{for} \quad t_n \leq t \leq t_{n+1}, \tag{S12}$$

where $t_{n+1} - t_n$ is the time interval between consecutive discrete responses. An integration model based on averaging of detected stimuli between consecutive responses can be performed as well, but no ambiguous differences in the results were observed.

Now the simulation results obtained by the discrete models and the continuous models are compared. Comparison between the discrete model and the continuous model for a typical set of simulations of the three phototaxis mechanisms (light-dependent angular turn, delay in photoresponse, up-down photoresponse inversion) is shown in Fig. S2. The discrete model and the continuous model provide qualitatively similar trajectories (top row of Fig. S2), albeit the discrete model displays discrete changes in both the signal and the turning response (middle and bottom row of Fig. S2). The transition in phototaxis behavior (i.e., from positive to negative phototaxis) is not influenced by the discreteness of the model. Hence the continuous model sufficiently captures the response observed in real biological cells with discrete flagellar beats.

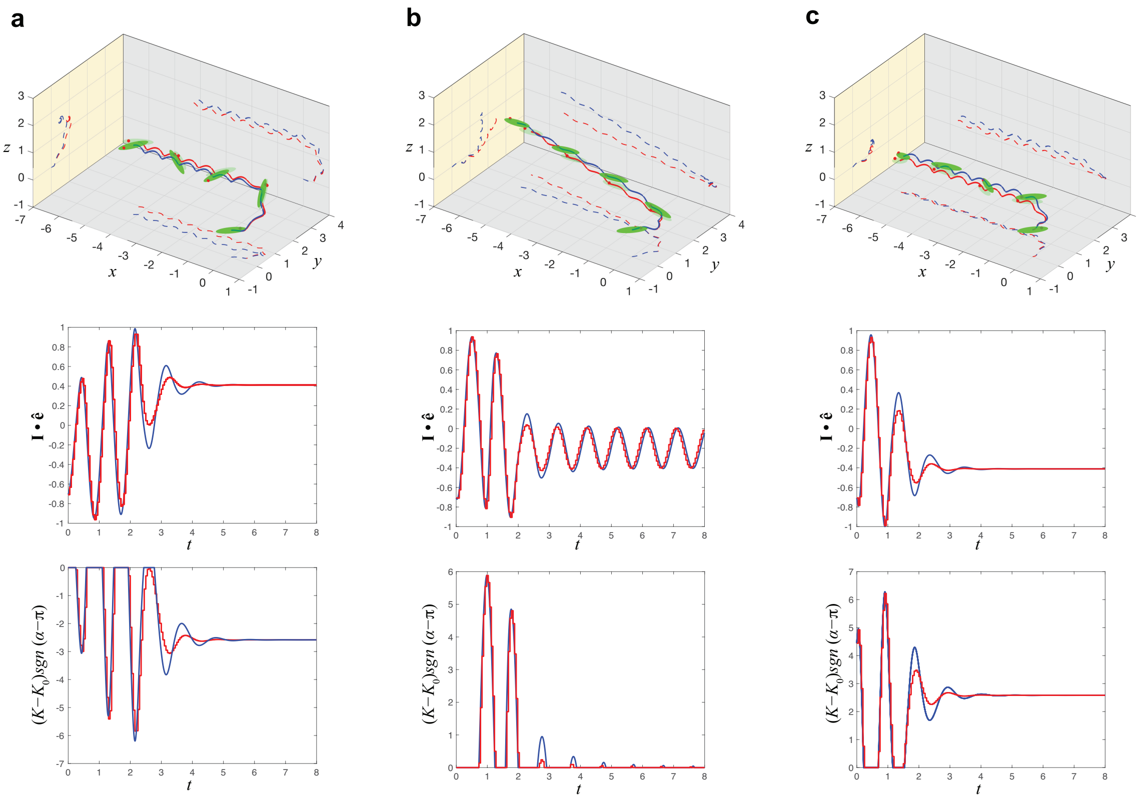


Figure S2: **Comparison of the three phototaxis mechanisms with a discrete or continuous response.** **(a)** Light-dependent angular turn mechanism. **(b)** Delay in photoresponse mechanism. **(c)** Up-down photoresponse inversion mechanism. The blue and red lines correspond to the results for the continuous model and the results for the discrete model, respectively.

#### 3.6 Attenuation of light penetrating into water

Here the attenuation of light penetrating into a water medium are compared. An LED with its glass cover hitting a water surface is considered, and the light intensity of the LED at different water depth was measured with a lux meter. The experimental data was then fitted with the modified Lambert's law and inverse square law [5]:

$$|\mathbf{I}| = I_0 c \frac{e^{-\beta|\mathbf{x}-\mathbf{x}_0|}}{|\mathbf{x}-\mathbf{x}_0|^{\gamma}} \tag{S13}$$

Here $I_0$ is the light intensity at a reference location $|\mathbf{x}_0| = 0$ cm (i.e., location of water surface from the center of LED). $c$, $\beta$ and $\gamma$ are coefficients accounting for optical aberration, absorption and path loss. The light intensity data was normalized with $I_0$. 3 sets of experiments were performed with different $I_0$ and the normalized data for all cases were found to be quantitatively similar (less than 5% differences for all data points). The experimental data was averaged and fit it with Eq. S13 (Fig. S3). The 95% confident interval of $c$, $\beta$ and $\gamma$ are given by $c = 0.79 \pm 0.08$, $\beta = 0.29 \pm 0.07$ and $\gamma = 0.074 \pm 0.012$.

#### 3.7 Photon count during positive phototaxis

In our experiments, the measured light intensity during positive phototaxis is $|\mathbf{I}| \sim 50$ lux. Assuming a luminous efficacy of 60 lm/W, such a light intensity amounts to 0.8 W/m$^2$. This result is consistent to numerous measurements in the literature [6]. Here the approximate amounts of photons is calculated that hit the photoreceptors when the photoreceptor is exposed to this light intensity. The total number of photons $N_{photon}$ illuminated on a Euglena's photoreceptor

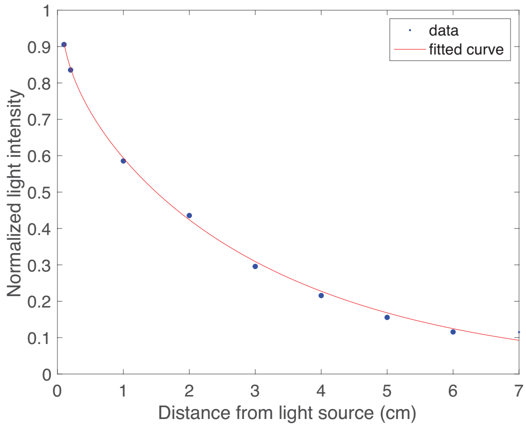


Figure S3: **Fitting of experimental data by modified Lambert's law and inverse square law given by Eq. S13.**

per second can be approximated by

$$N_{photon} = |\mathbf{I}| \times A_{receptor}/E. \tag{S14}$$

Here, $A = 1 \times 10^{-12}$ m$^2$ is the projected area of the photoreceptor [7]. $E_{photon}$ is the energy carried by a photon. Here a blue light with wavelength $\lambda = 4.7 \times 10^{-7}$ m, $E_{photon} = hc/\lambda$ is considered, and where $h = 6.626 \times 10^{-34}$ j·s is the Planck's constant and $c = 3 \times 10^8$ m/s is the speed of light. Hence $N_{photon} \sim 2 \times 10^6$.

##### 3.8 Extended analysis for phototaxis with two light sources

The current work mainly focuses on the photoresponses towards a single light source. Here now a potential extension of the current model is discussed to include multiple light sources. This analysis is motivated by previous experiments [8].

First, the case of two perpendicular light beams in the x-y plane is considered (Fig. S4a). The two light beams have the same high light intensities and that stay constant over the considered distances. The negative phototaxis model then shows that the cell eventually swims in a direction of the resulting vector of both light sources (i.e., $\sim 45°$ with respect to the two light sources), in agreement with the previously reported experiments [8].

Second, the the case of two opposite light beams in the x-y plane at equally low light intensities is considered (Fig. S4b), where the light intensities decay according to Eq. (S13). The photoresponse inversion model shows that the cell will eventually swim towards either one of the light sources. Initially when the cell is situated in the middle of the two light sources, the cell detects equal strength of both opposite light sources which are low relative to the noises due to rotational diffusion of the cells. Hence the cell has no specific preference in the swimming direction initially. However, due to the random motion of the cell, it will eventually receives a slightly larger signal from one of the light sources and bias its swimming path towards it. Yet, this initial model investigation for positive phototaxis under two light sources do not replicate a clear bimodal distributions of cell orientation as reported in previous experiments [8], which will require a deeper analysis in both modeling and experiments as the reason could be in either one, and which is beyond the present work.

##### 3.9 Model parameters for simulations

Here, the model parameters are summarized that have been used for the simulations shown in main text Fig. 2, Fig. 3 and Fig. 7. For all simulations, the system was nondimensionalized using

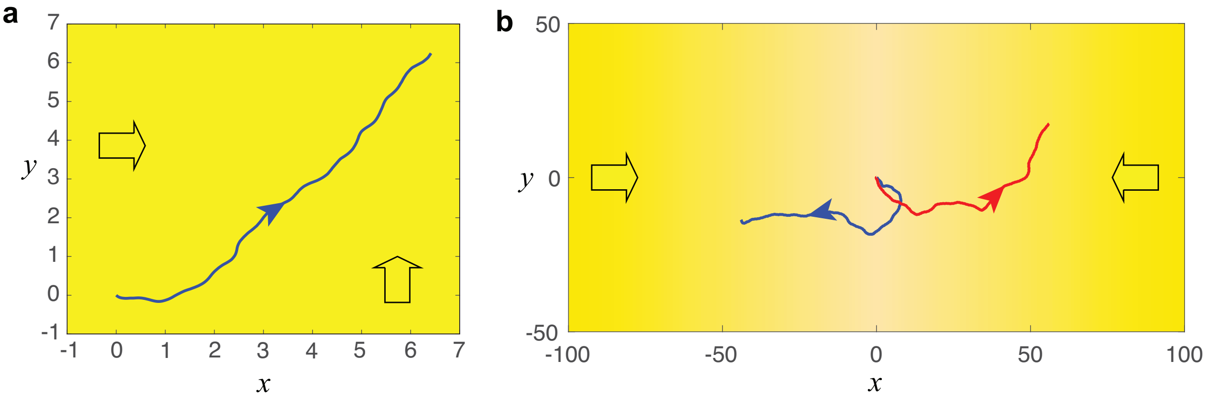


Figure S4: **The model recapitulate the phototactic responses of cells under two light sources from two different directions. (a)** When the cell is subject to two strong light sources, it performs negative phototaxis and swim in the direction of the resulting light vector of both light sources. $\mathbf{I}_1 = 2\hat{\mathbf{x}}$ and $\mathbf{I}_2 = 2\hat{\mathbf{y}}$. $v = 1$, $\alpha = 3\pi/2$, $K_0 = 0.3\omega$, $K_a = 0$, $K_d = 0.03\omega$, $D_r = 0.02$, $D_t = 8 \times 10^{-8}$, $\omega = 2\pi$, $I_\epsilon = 0.025$; **(b)** When the cell is subject to two weak light sources with a signal decay over distances, the cell swims towards either one of the light sources. The light intensity $|\mathbf{I}_1|$ and $|\mathbf{I}_2|$ is given by Eq. (S13). The parameters were chosen to be: $c = 0.79$, $\beta = 0.29$, $\gamma = 0.074$, $I_0 = 0.1$, $x_0 = -200$ for $\mathbf{I}_1$ and $x_0 = 200$ for $\mathbf{I}_2$.

the body length $\ell$ as the length scale and $1/\omega$ as the time scale, where $\omega$ is the rolling frequency. The swimming velocity $v$ was chosen to be consistent with the experimental measured swimming velocity of the cell.

**Simulations in main text Fig. 2:**

For the simulations shown in main text Fig.2 (blue solid lines), the simulations were performed using the light-dependent angular turn mechanism, but we note that all three mechanisms yield qualitative similar results. The coupling constant $k_d$ was chosen to fit the experimental measured statistics. The non-dimensional parameters are summarized as follows:

**Negative phototaxis (Fig. 2b):** $\mathbf{I} = -2\hat{\mathbf{y}}$, $v = 1$, $\alpha$ is selected randomly from $(\epsilon + \pi/2, 2\pi - \epsilon/2)$, $K_0 = 0.3\omega$, $K_a = 0$, $K_d = 0.03\omega$, $D_r = 0.02$, $D_t = 8 \times 10^{-8}$, $\omega = 2\pi$, $I_\epsilon = 0.025$;

**Positive phototaxis (Fig. 2d):** $\mathbf{I} = -0.05\hat{\mathbf{y}}$, $v = 1$, $\alpha$ is selected randomly from $(\epsilon - \pi/2, \pi - \epsilon/2)$, $K_0 = 0.3\omega$, $K_a = 0$, $K_d = 0.2\omega$, $D_r = 0.02$, $D_t = 8 \times 10^{-8}$, $\omega = 2\pi$, $I_\epsilon = 0.025$;

**Simulations in main text Fig. 3:**

For the simulations shown in main text Fig. 3, a typical simulation is shown for negative phototaxis and simulations for positive phototaxis exhibited by the three mechanisms according to main text Eq.(1)-(3). No rotational,translational and sensory noises were added in the simulations. The parameters are summarized as follows:

**Negative phototaxis (Fig. 3a, Movie S4):** $\mathbf{I} = 1\hat{\mathbf{x}}$, $v = 1$, $\alpha = 5\pi/4$, $K_0 = 0.3\omega$, $K_a = 0$, $K_d = 1$, $\omega = 2\pi$;

**Positive phototaxis: light-dependent angular turn (Fig. 3b, Movie S5):** $\mathbf{I} = 1\hat{\mathbf{x}}$, $v = 1$, $\alpha = \pi/4$, $K_0 = 0.3$, $K_a = 0$, $K_d = 1$, $\omega = 2\pi$;

**Positive phototaxis: delay in photoresponse (Fig. 3c, Movie S6):** $\mathbf{I} = 1\hat{\mathbf{x}}$, $v = 1$, $\alpha = 5\pi/4$, $K_0 = 0.3$, $K_a = 0$, $K_d = 1$, $t_d = 0.5$, $\omega = 2\pi$;

**Positive phototaxis: inversion in photoresponse (Fig. 3d, Movie S7):** $\mathbf{I} = 1\hat{\mathbf{x}}$, $v = 1$, $\alpha = 5\pi/4$, $K_0 = 0.3$, $K_a = 0$, $K_d = 1$, $\omega = 2\pi$ $\Delta = 0.2$.

**Simulations in main text Fig. 7:**

For the simulations shown in main text Fig. 7c-f (Supplementary Movies S12, S13), positive phototaxis was considered for the three mechanisms under two types of light stimulus. i.e., parallel light field and light gradient. The simulations were performed using main text Eq.(1)-(6). The parallel light field is given by $\mathbf{I} = 0.2\hat{\mathbf{x}}$. The light gradient in $\hat{\mathbf{x}}$-direction is given by combining Lambert's law and inverse square law, $\mathbf{I} = |\mathbf{I}|\,\hat{\mathbf{x}}$, where the light intensity $|\mathbf{I}|$ is given by Eq. (S13). The parameters were chosen to be: $x_0 = 200$, $c = 0.79$, $\beta = 0.29$, $\gamma = 0.074$, $I_0 = 0.1$, which is consistent with the decay of light in pool water [5] (See Supplementary Section 3.6).

For all mechanisms, the following common parameters were chosen: $v = 1$, $K_0 = 0.3$, $K_a = 0$, $K_d = 0.2\omega$, $D_r = 0.02$, $D_t = 8 \times 10^{-8}$, $I_\epsilon = 0.025$, $\omega = 2\pi$. For each mechanism, the following parameters were chosen: light-dependent angular turn: $\alpha = \pi/2$; delay in photoresponse: $\alpha = 3\pi/2$, $t_d = 0.5$; inversion in photoresponse: $\alpha = 3\pi/2$, $\Delta = 0.2$.

### 4 Supplementary experiments and analysis

#### 4.1 Validation of cell samples for positive phototaxis

The response in positive phototaxis is slow and noisy compared to negative phototaxis (main text Fig. 2), where noticeable responses are observed in the order of seconds for negative phototaxis and in the order of minutes for positive phototaxis. Cells in different samples can have slightly different light intensity thresholds for triggering positive phototaxis. Moreover, cells tend to show weak responses to light stimuli at low light intensities when they had been adapted to the ambient light at medium light intensities that were generally used for keeping the cells. Therefore, in order to obtain robust and consistent experimental results for positive phototaxis experiments, it needed to be validated that cell cultures can display robust positive phototaxis behaviors.

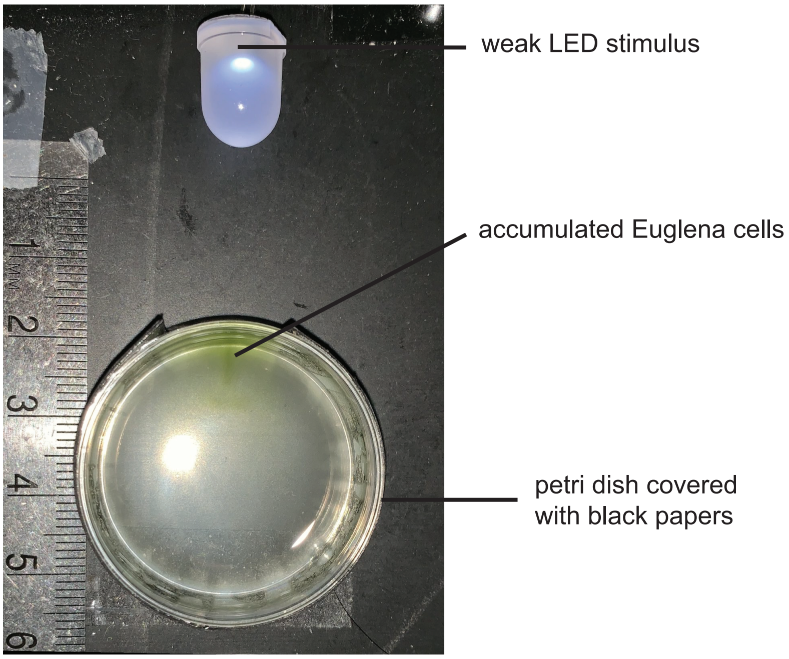


Figure S5: **The positive phototaxis of a Euglena cell sample towards the weak LED stimulus was tested to ensure that the cells are responsive to the LED at low light intensity (~ 50 lux).** The greenish color in the petri dish denotes the accumulated cells due to positive phototaxis. The experiment was conducted inside a black box that block all the ambient light. The cover of the box was opened at 10 minutes after the LED stimulus was introduced to observe the result. The unit numbers on the ruler are in cm.

To this end, the Euglena cells were first kept in dark for 24 hours such that they were fully adapted to darkness. The cells were then placed inside a petri dish covered with black papers to avoid any reflection of light inside the dish, except for a small opening that allowed light to enter the dish. A weak LED stimulus was introduced to the cells and the cells slowly accumulated towards the small opening (Fig. S5), displaying a greenish color given by the accumulated Euglena cells after 10 minutes (other regions of the petri dish remained mostly colorless). This test ensured that the cells exhibited positive phototaxis towards the light intensity level we considered.

These responsive cells in the region of accumulation were then collected and transferred to a flow chamber for further experiments. The same amount of light stimulus was introduced to the cells in the flow chamber via a side LED, with red observing light from the microscope. The LED was placed in the same distance from the cells compared to the petri dish experiment. This validation procedure ensured that the investigated cells were positively phototactic towards the light stimulus. Note also that some cell cultures showed robust positive phototaxis without the need of keeping the cells in dark for 24 hours. The validation procedure was performed for robustness of results. The change in photoresponse of cells under long time adaptation (i.e., in the order of hours and days) to different light intensity levels are beyond the scope of the current work.

#### 4.2 Tracking the phototaxis statistics of Euglena cells

The phototaxis statistics of Euglena cells (main text Fig. 2c,d & 7a,b) were obtained by tracking a few hundreds to a few thousands of Euglena cells using an automatic tracking program developed in a previous work [9]. The cells were tracked for 5 minutes before and after introduction of light stimuli, where each cell was tracked for at least 3 seconds. The center position, orientation, and velocity of the cells were tracked.

It was noted that all three phototaxis mechanisms can effectively capture the orientation statistics of Euglena cells under positive phototaxis (Fig. S6). No difference in the overall orientation statistics for the three models was observed, as can be seen from the overlapping lines in Fig. S6.

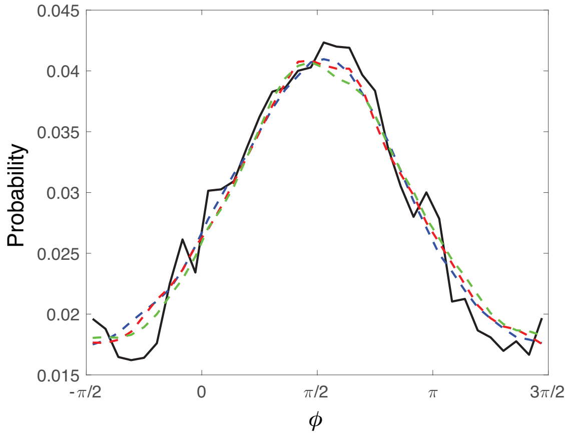


Figure S6: **Comparison of experimental statistics of cell orientation under positive phototaxis with the model predictions given by the three mechanisms.** The black solid line denotes the experimental data ($N = 406$ cells). The blue, red, green dashed lines represent the cell orientation statistics of light-dependent angular turn mechanism, delay in photoresponse mechanism, up-down photoresponse inversion mechanism, respectively. The results are obtained from 1000 runs of Monte-Carlo type simulations.

For experiments that required a longer time of tracking, in particular, the measurement of the mean free paths of the cell under darkness and positive phototaxis, the positions of 50 cells were manually tracked for each case, where each cell was tracked at a sampling rate of 1 s for at least 8 seconds. Fig. S7 depicts some example trajectories for cells during negative photoaxis and positive phototaxis. The manual tracking allowed to obtain data with long enough swimming paths to measure the mean free paths of the cells. The data was used to calculate the displacement direction correlation function (main text Fig. 7a) [10]:

$$C_\tau(t) = \langle \mathbf{d}(0,\tau) \cdot \mathbf{d}(t, t+\tau) \rangle, \tag{S15}$$

where $\mathbf{d}(t, t+\tau)$ is the direction of the unit vector obtained by joining the body center of the cells at time $t$ and time $t+\tau$.

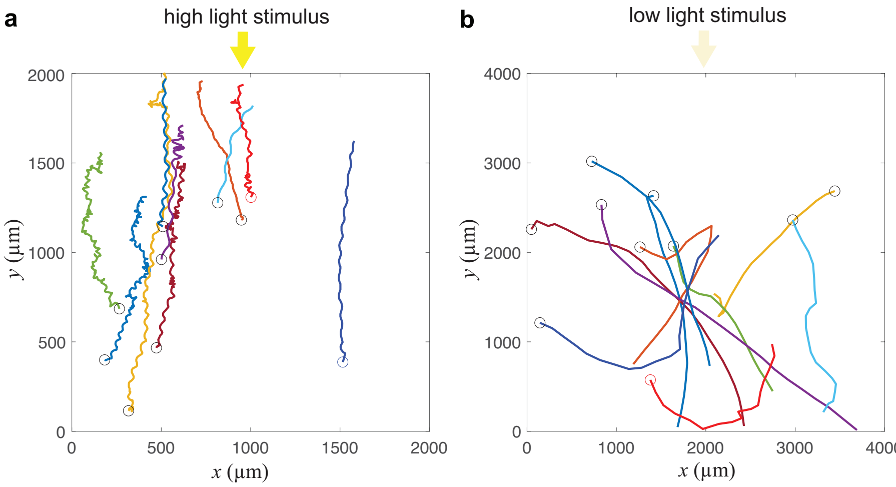


Figure S6: **Example trajectories for cells undergoing negative phototaxis and positive phototaxis.** **(a)** Cells exhibit directional swimming towards light stimulus under negative phototaxis. **(b)** Cells do not display obvious directional swimming under positive phototaxis.

In main text Fig. 7a, it is shown that Euglena cells exhibiting positive phototaxis display a smaller $C_\tau$ over times compared to cells in darkness, indicating that the cells exhibitng positive phototaxis have a shorter mean free paths ($n = 50$, $p < 0.01$ over 8 s of data). It was also shown that the variance of the reorientation rate for the cells exhibiting positive phototaxis is larger than for the cells in darkness, indicating that the positively phototactic cells reorient more frequently. For completeness, the displacement direction correlation and the reorientation rate of negatively phototactic cells was also plotted in Fig. S8. The correlation function remains large over a relatively longer time, and the reorientation rate has a narrower distribution, which shows that the negatively phototactic cells swim in the same direction with relatively small adjustments of their paths.

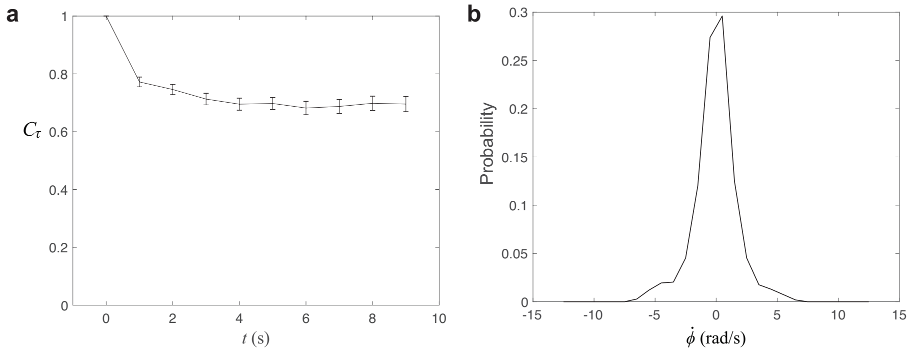


Figure S8: **Statistics of negative phototactic cells display directed swimming.** **(a)** Displacement correlation function. **(b)** Distribution of reorientation rate.

For all tracking experiments, the tracking of cells was repeated in at least three different cell cultures on at least three different days and different times of the day (9am to 9pm), and consistent results were obtained in all our tracking data. This confirmed the robustness of our experimental tracking results.

##### 4.3 Measuring the paw angle $\alpha$ of Euglena cells

For experiments shown in main text Fig. 4, the paw angle $\alpha$ of freely swimming cells was measured under various conditions, i.e., (i) helical swimming in darkness ($\sim$ 0 lux), (ii) positive phototaxis under directional stimulus ($\sim$ 50 lux), (iii) helical swimming under medium background ($\sim$ 100 lux), (iv) negative phototaxis under high stimulus ($>$ 2000 lux). In our previous work [2], it had been shown that the frequencies of body rolling and helical swimming are coupled, identical, and phase locked, where there is a fixed phase relation between $\psi$, $\phi$ and $\theta$ for a given $\alpha$ (see main text Fig. 4a,b). Therefore, the paw angle $\alpha$ can be obtained by measuring the intercept of the $\psi$ and $\theta$ plot (main text Fig. 4b).

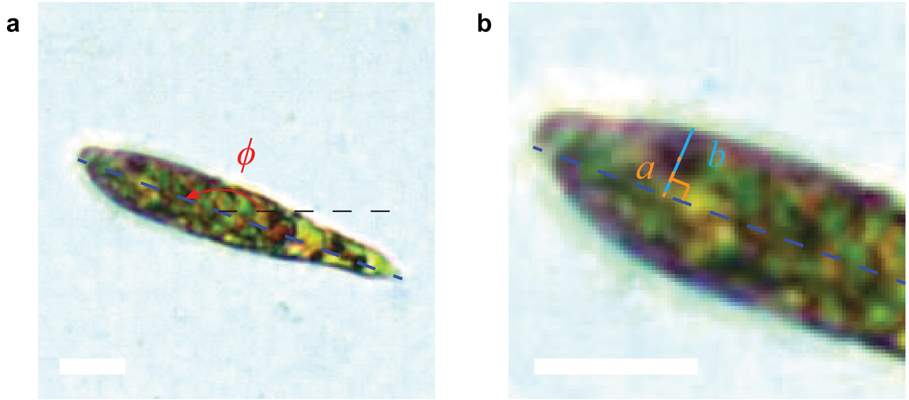


Figure S9: **Tracking cell orientation and eyespot of the Euglena cells determines the paw angle for different taxis behavior. (a)** Tracking of cell orientation. **(b)** Tracking of eyespot. The blue dashed line denotes the centerline of the cell body. The red dashed line denotes the location of the eyespot center measured from the body centerline. The orange dashed line denotes the farthest location where the eyespot can reach. Scale bars: 5 $\mu$m

The orientation $\phi$ of the swimming cells was tracked under each condition (Fig. S9). The roll angle $\psi$ was also tracked using the eyespot as reference point when observed from the top, i.e., $-\hat{\mathbf{z}}$, $\sin(\psi) = b/a$, where $a$ is defined to be positive when the eyespot is on the left side of the body centerline and $b$ is always positive. The cell orientation $\phi$ was then mapped to the orientation phase $\theta$ according to the oscillation in orientation of the helical swimming motions (main text Fig. 4a).

A total of 41 cells were tracked for at least 3 rolling cycles to obtain the $\alpha$ of different behaviors via fitting of the experimental tracking data (main text Fig. 4d). The results showed that there is no significant difference in the experimentally measured $\alpha$ between cells exhibiting positive phototaxis and negative phototaxis. This rules out the light-dependent angular turn mechanism.

##### 4.4 Tracking flagellar beats of Euglena cells using micropipette aspiration

For experiments shown in main text Fig. 5 and Fig. 6, micropipette aspiration was used to capture and fix a Euglena cell in place, enabling us to keep the cells in focus for long times and observe the response of flagellar patterns upon changes in light intensity. In particular, four light conditions at different light intensities were considered, namely darkness ($\sim$ 0 lux), low light ($\sim$ 50 lux), medium light ($\sim$ 100 lux) and high light($\sim$ 5000 lux) to mimic conditions of transition in positive phototaxis and negative phototaxis.

The cells were imaged at 400 fps. The flagellar beat patterns of the cells were tracked in ImageJ. The observed beat patterns were quantified with the angle $\lambda$ between the cell tip and the farthest material point on the flagellum within a beat cycle (main text Fig. 6a). Two main beat patterns corresponding to helical swimming and turning behavior were observed (main text Fig. 6a), where the two beat patterns featured distinct beat angles. The beat pattern for helical swimming has a beat angle of $\lambda \sim 0$ or $\lambda \sim \pi$ (depending on how cell was oriented) and the beat patterns for turning behavior has a beat angle of $\lambda \sim \pi/2$ (the flagellum is mostly situated in front of the cell). It was noted that different cells may have slightly different $\lambda$ for their swimming and turning beat patterns.

The change in beat angle $\lambda$ was used to define a beat switching event. For each cell we considered, the average $\lambda$ for the swimming and turning beat patterns was calculated. These two average values of $\lambda$ were then used as references to define whether a beat switching occurs. If a cell exhibiting a helical swimming beat pattern changed its beat pattern, and the $\lambda$ of the updated beat pattern was different from the average $\lambda$ of helical swimming beat patterns by a certain threshold (this threshold was set to be $\pi/6$, see main text Fig. 3a and main text Fig. 6c for typical difference in $\lambda$ between the two beat states), then this event was regarded as a beat switching. Using slightly different thresholds will yield similar results. The same rule was used to account for the beat switching from the turning beat patterns. The switching between the swimming beat patterns and the turning beat patterns was continuous and involved a small number of transient beats. These transient beats, which had $\lambda$ values not covered in the range of the average $\lambda \pm \pi/6$ of the two beat patterns, were neglected in this study.

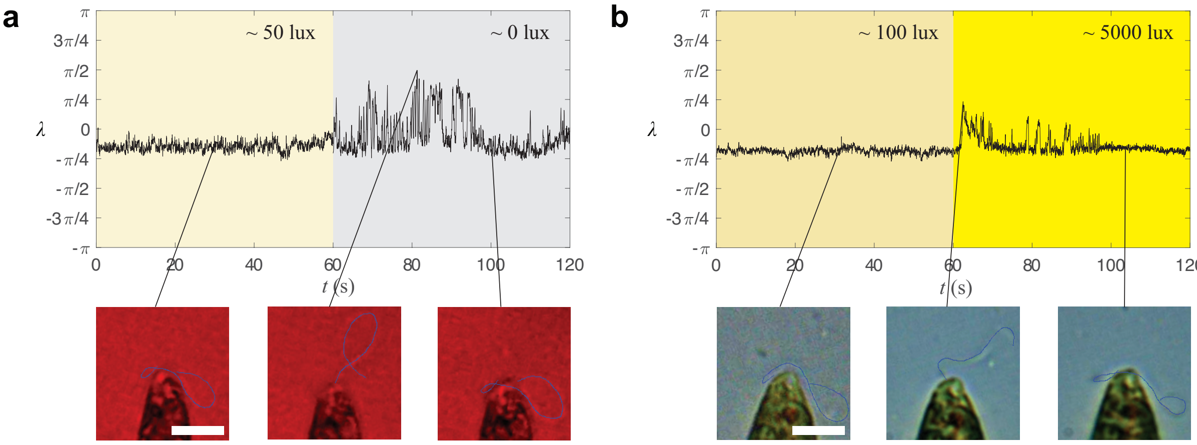


Figure S10: **The beat patterns of Euglena stochastically switched back and forth between the beat patterns for helical swimming and turning when the light conditions correspond to the responsive regime of the cells, as indicated by the fluctuation in the beat angle $\lambda$.** **(a)** Under positive phototaxis condition, when a weak LED stimulus ($\sim$ 50 lux) is off and the light condition becomes darkness ($\sim$ 0 lux), the cell changed its beat state from stable beat patterns for helical swimming to a state where the beat patterns switched stochastically between the beat patterns for helical swimming and turning. **(b)** Under negative phototaxis condition, when a cell experience a step-up in light intensities from medium light ($\sim$ 100 lux) to high light ($\sim$ 5000 lux), the cell changed its beat state from stable beat patterns for helical swimming to a state where the beat patterns switched stochastically between the beat patterns for helical swimming and turning, with $\lambda$ decaying almost monotonically over time. The bottom panels show example snapshots of flagellar beat patterns at different responsive regimes. Scale bars: 5 $\mu$m

The transient and adaptation behavior of flagellar response was observed by periodically turning the stimulus on and off. Under negative phototaxis condition, the light condition was alternated between medium light and high light, while under positive phototaxis condition, the light condition was alternated between darkness and low light. The on-period and the off-period of the light stimulus last for at least half a minute. In main text Fig. 6c, the beat angle $\lambda$ is presented over times before and after light changes. It was found that the responses between the negative and positive phototaxis conditions agreed with the inversion of photo-response mechanism. Under negative phototaxis condition, a stronger response (transition from swimming beat to turning beat) was observed when the stimulus was on, while under positive phototaxis condition, a stronger response was observed when the stimulus was off.

The adaptation of the cell response over longer times was also considered (Fig. S10). After the light condition was changed to the regime where the response was the strongest (i.e., on-period for negative phototaxis and off-period for positive phototaxis), the beat pattern stochastically switched back and forth between the beat pattern for helical swimming and turning for a while (typically around $1 - 3$ mins) before the cells fully adapted to the new light condition and only exhibited beat patterns for helical swimming. Compared to the case of positive phototaxis condition, the beat angle $\lambda$ of cells under negative phototaxis condition decayed more monotonically over times. It was noted that the response for negative phototaxis was typically more robust than positive phototaxis with predictable results.

Finally, it was noted that a small number of captured cells ($< 10\%$) largely deformed their bodies due to suction flows from the micropipette (most cells only deformed their bodies slightly). The abnormal behaviors of these deformed cells were out of scope in our current study and were therefore not included in the current analysis.
